# TXTL-Powered K1F Internal Capsid Protein Engineering for Specific, Orthogonal and Rapid Phage-based Pathogen Detection

**DOI:** 10.1101/2024.05.06.592667

**Authors:** Joseph P. Wheatley, Sahan B. W. Liyanagedera, Tamás Fehér, Antonia P. Sagona, Vishwesh Kulkarni

## Abstract

The internal capsid proteins that reside within phage of the Podoviridae family hold high potential for being used as sensitive and reliable diagnostic tools. The concealed nature of the capsid interior ensures that any encapsulated signal or signal generating enzyme, e.g., fused to an internal capsid protein, is suppressed whilst the phage is unaccompanied by its host. Furthermore, the only naturally occurring mechanism for releasing the internal capsid proteins, and therefore exposing their amalgamated signal/enzyme, is for them to be passed through the tail and subsequently ejected out of the phage, a post-adsorption phenomenon which occurs when the host is present, thus presenting a precise model for signal/enzyme release only upon pathogen presence. Here, a small N terminal subunit of the NanoLuc luciferase is fused and incorporated into the K1F internal capsid structure using a simple, non-genomic method. This internalised subunit is exposed to the test solution containing its C terminal counterpart (natural complementation immediately forms the full NanoLuc enzyme) and substrate furimazine in an inducible manner which mimics the presence of the K1F host, E. coli K1 thereby presenting a novel method for rapidly detecting this disease causing pathogen. Finally, it is expected that by building upon this internal capsid protein engineering approach, which completely bypasses the time-inducing processes of intracellular nucleic acid transcription and translation, an unprecedentedly rapid detection device can be developed for an array of bacterial pathogens.

## 1. Introduction

The increasing incidence of bacterial infections worldwide, alongside the often uninformed overuse of antibiotics, have harboured an urgent need for rapid pathogen detection. A fast and specific diagnosis would allow for early, targeted therapy to be administered and for antibiotics to be spared for necessary cases only, subsequently leading to a decrease in medical burden and antimicrobial resistance (AMR) [26].

The traditional culture-based method has been the gold standard of bacterial identification for decades. However, due to the time-consuming (> 24 hours) and laborious nature of bacterial culturing, many alternative detection methods have been established and continue to be developed. Existing alternative methodologies/technologies for bacterial detection include: phage-based signal amplification, enzyme linked immunosorbent assay (ELISA), gold nanoparticle (AuNP) aggregation, polymerase chain reaction (PCR), nucleic acid sequence-based amplification (NASBA), loop-mediated isothermal amplification (LAMP) and recombinase polymerase amplification (RPA). A comparative display of recent *E. coli* detection systems built utilising these methods and their capabilities is shown in Table 1. Furthermore, whilst these detection methods each have their own advantages and disadvantages which have been discussed in previous reviews [11,12], they are all uniform in the fact that they require additional technical expertise or equipment in order to complete their respective diagnostic tests. In this instance, technical expertise or equipment is defined as necessitating at least basic laboratory skills and/or equipment and therefore rendering the test unusable in a standard General Practice (GP) or test-at-home scenario. There are also numerous commercial lateral flow test (LFT) *E. coli* antigen detection systems available [13,14]. However, before a diagnosis can be given they require the user to carry out an enrichment step on their sample to cultivate the bacteria, meaning that only next-day results (16+ hours) are achievable.

**Table 1.**
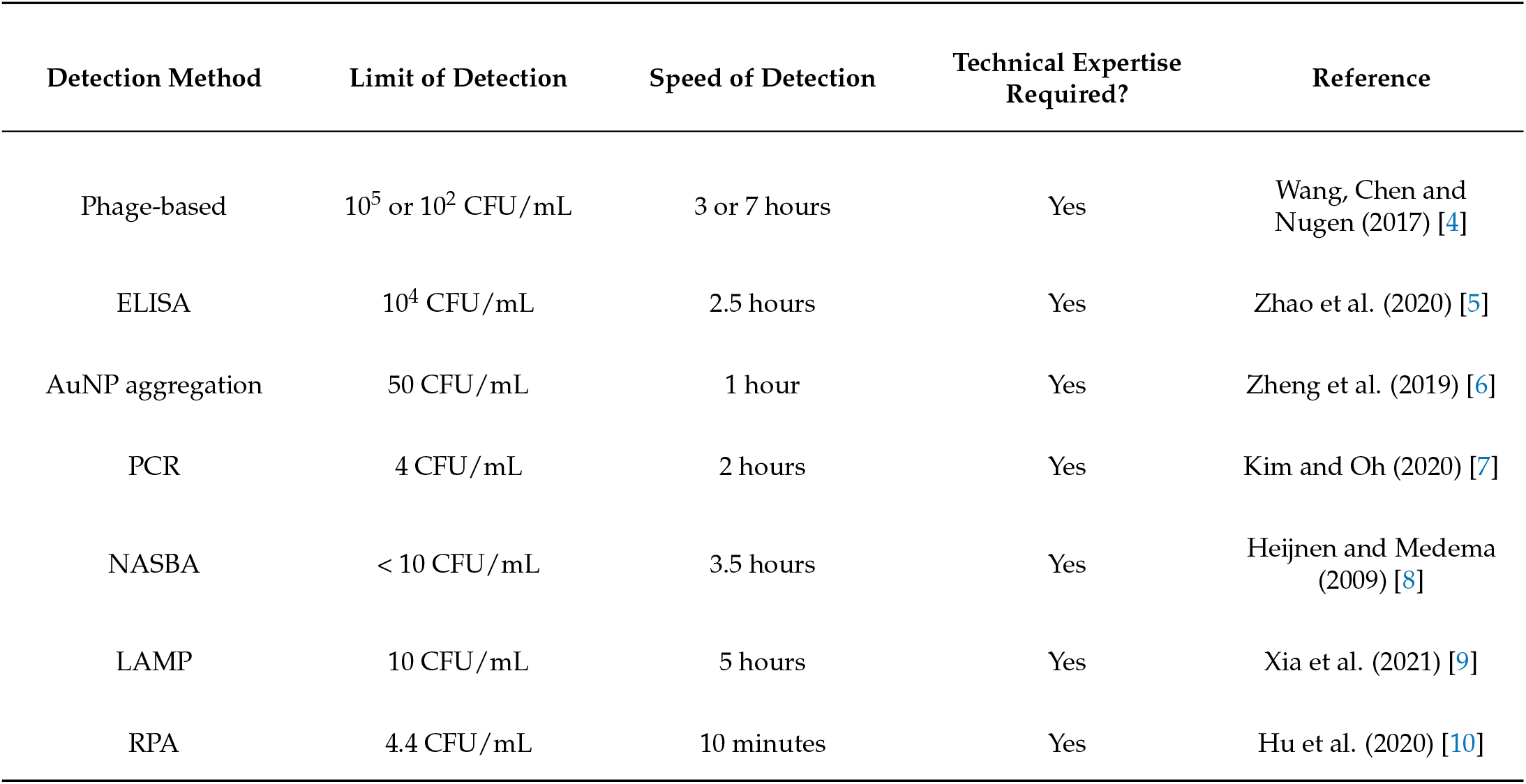
Bacterial detection method case-study comparison.

| Detection Method | Limit of Detection | Speed of Detection | Technical Expertise Required? | Reference |
| --- | --- | --- | --- | --- |
| Phage-based | 10<sup>5</sup> or 10<sup>2</sup> CFU/mL | 3 or 7 hours | Yes | Wang, Chen and Nugen (2017) [4] |
| ELISA | 10<sup>4</sup> CFU/mL | 2.5 hours | Yes | Zhao et al. (2020) [5] |
| AuNP aggregation | 50 CFU/mL | 1 hour | Yes | Zheng et al. (2019) [6] |
| PCR | 4 CFU/mL | 2 hours | Yes | Kim and Oh (2020) [7] |
| NASBA | < 10 CFU/mL | 3.5 hours | Yes | Heijnen and Medema (2009) [8] |
| LAMP | 10 CFU/mL | 5 hours | Yes | Xia et al. (2021) [9] |
| RPA | 4.4 CFU/mL | 10 minutes | Yes | Hu et al. (2020) [10] |

This publication includes results based on cell-free TXTL phage assembly. In what was a monumental achievement at the time, the first demonstration of cell-free infectious phage synthesis was a decade ago [15], where phage T7 and *ϕ*X174 were used as model organisms. Since then, there have been numerous innovations within the TXTL phage synthesis space [16].

In the work presented in this publication, a proof-of-concept phage-based bacterial detection system is hypothesised, designed and tested. It is envisioned the technology developed here could produce: 1). a diagnosis in minutes, rather than hours; 2). an orthogonal and adaptable detection approach capable of simultaneously screening for multiple pathogens; 3). an end-product that requires no technical expertise or equipment; and 4). the first truly rapid test-at-home device for multiple bacterial pathogens.

The phage used in this study to demonstrate this novel detection model is K1F. The T7-like phage K1F is of particular interest because of its cognate host, *E. coli* K1, a gramnegative pathogen responsible for a wide range of potentially fatal diseases in humans, including sepsis, neonatal meningitis, urinary tract infections and inflammatory bowel syndrome [17].

## 2. Materials and Methods

### 2.1 Plasmid construction

Plasmid pBEST-OR2-OR1-Pr-UTR1-deGFP-T500 (Addgene 40019) was the kind gift of Vincent Noireaux (The University of Minnesota), obtained via Addgene. The three expression plasmids used, pBEST-OR2-OR1-Pr-UTR1-g6.7::NtNL-T500, pBEST-OR2-OR1-Pr-UTR1-g14::NtNL-T500 and pBEST-OR2-OR1-Pr-UTR1-CtNL-T500 were constructed by digesting synthetic gene blocks ordered from Integrated DNA Technologies (Leuven, Belgium) with NcoI and XhoI and ligating to the corresponding restriction sites of the pBEST-OR2-OR1-Pr-UTR1-deGFP-T500 plasmid, thereby replacing deGFP with the desired gene sequence. For pBEST-OR2-OR1-Pr-UTR1-g6.7::NtNL-T500 and pBEST-OR2-OR1-Pr-UTR1-g14::NtNL-T500 the synthetic gene blocks comprised the K1F internal capsid protein gene (g6.7 or g14) fused to the N-terminus of the NanoLuc gene by a (Gly-Gly-Gly-Gly-Ser)2 linker and for pBEST-OR2-OR1-Pr-UTR1-CtNL-T500 the synthetic gene block comprised the C-terminus of the NanoLuc gene (for exact DNA sequences, see Table S1).

#### 2.1.1 Measurement of bioluminescence

Bioluminescence was measured using a FLUOstar Omega microplate reader (BMG Labtech, Aylesbury, England) and the assay was carried out using the Nano-Glo® Luciferase assay system and associated protocol (Promega, Madison, USA). Samples were mixed with an equal volume of the Nano-Glo® assay reagent and 20 *µ*L of this mix was subsequently pipetted into the wells of a Nunc™ 384-well microplate (Thermo Fisher Scientific, Waltham, USA). The relative luminescence was measured within one hour of mixing the assay reagent and samples together. If combining the N- and C-terminal NanoLuc sub-units prior to the luminescence assay, unless otherwise specified the two sub-units were mixed and incubated at room temperature for one hour before adding the assay reagent to allow for spontaneous complementation.

#### 2.1.2 Bacteriophage propagation

A high K1F phage titer was achieved by propagating the phage over a series of infection cycles in cultures with increasing *E. coli* EV36 concentration - starting with a culture with an *OD*_600_ value of 0.2, followed by 0.8 and finally 1.2. Immediately after bacterial clearance was observed, each propagation culture was centrifuged at 3220 g for 15 minutes at 4 °C and the supernatent was obtained. After the clearance and centrifugation of the final culture (*OD*_600_ 1.2), the supernatent was passed through a filtration unit to remove the majority of any bacterial remains still present. At this stage, 1 *µ*g/mL of both DNase I and RNase I (New England Biolabs, Ipswich, USA) was added and incubated with the phage at room temperature for 1 hour to digest any non-phage nucleic acid present in the mix. Next, 0.2 M NaCl was added and incubated on ice for 1 hour to facilitate the release of phage particles from bacterial membranes. Following on from this was a centrifugation at 5000 g for 45 minutes at 4 °C. The supernatant was recovered and incubated with 10% w/v PEG8000 overnight at 4 °C to precipitate phage particles. The following day, the PEG-phage solution was centrifuged for 25,000 g for 1 hour at 4 °C and the pellet was resuspended in SM buffer.

#### 2.1.3 Bacteriophage purification

A CsCl density gradient was prepared by mixing varying amounts of CsCl with water to produce three densities of 1.7, 1.5 and 1.3 g/ml. CsCl was then added to the SM buffer-phage solution to achieve a density of 1.3 g/ml. Using a manual pipette, the CsCl solutions were slowly added to a centrifuge tube in equal volumes starting with the heaviest. Once the CsCl-phage tube was prepared, it was placed in a SW28.1 rotor and centrifuged in a Beckman L-90K ultracentrifuge (Beckman Coulter, Pasadena, U.S.A) at 125,000 g for 20 hours at 4 °C. The resulting blue/grey band was extracted and transferred into a Slide-A-Lyzer™ 10k MWCO cassette (Thermo Fisher Scientific, Waltham, USA) which was then dialysed overnight (approximately for 16 hours) in SM buffer. Following on from dialysis, the purified phage were stored at 4 °C.

#### 2.1.4. Bacteriophage DNA isolation

CsCl purified phage were used for DNA isolation. An equal volume of Tris-saturated phenol (pH 8.0) was added to the phage suspension, vortexed and left at room temperature for 10 minutes. The tube was spun down at 13,000 g for 10 minutes at 4 °C in a 38 Rotanta 46R centrifuge (Hettich, Tuttlingen, Germany). The aqueous phase containing the phage at the top was extracted and added to a new tube. An equal volume of phenol:chloroform (1:1) was added to the extracted phage and vortexed. The tube was then centrifuged again at 13,000 g for 10 minutes at 4 °C. The top layer was extracted again and an equal volume of phenol:chloroform:isoamylalcohol (25:24:1) was added to the tube, vortexed and centrifuged again. 1/10th volume 7.5 M ammonium acetate (pH 8.0) was added together with 2 volumes of ice-cold isopropanol and left overnight at -20 °C for precipitation. The next day, the tubes were centrifuged at 3220 g for 15 minutes at 4 °C (Avanti J-25/JA 25.5 rotor). The supernatant was removed and the pellet was washed with ice-cold ethanol. Finally, the pellet was air-dried and then resuspended in molecular grade water.

#### 2.1.5 Plaque assay

LB agar plates were overlain with 4 mL of top agar containing 400 *µ*L of *E. coli* EV36 cells (*OD*_600_ = 0.2) and 100 *µ*L of serially diluted phage. After solidification, plates were incubated overnight at 37 °C. The following day, the plaque numbers were counted and subsequently used to calculate phage titers in PFU/mL. The titer calculation is as follows:

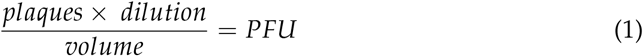

### 2.2 Electron microscopy analysis of cell-free TXTL phage synthesis

10 *µ*L drops of TXTL K1F assembly reaction from different time intervals were applied to the centre of the mesh and were incubated for 1 minute. The samples were removed and the mesh washed twice with 10 *µ*L drops of water and finally negatively stained with 10 *µ*L 2% uranyl acetate for 1 minute. Images were acquired using the Jeol 2100 transmission electron microscope (Jeol, Tokyo, Japan).

## 3. Results

### 3.1 Strategy

Given the high disease burden of bacterial infections and the serious impending threat posed by antimicrobial resistance, the development of a truly rapid and reliable bacterial detection system has been one of the great ambitions of modern day phage biology. Whereas the results produced in previous publications showcase diagnostic phage that can detect their host in a matter of hours [1,4,18], the novel approach demonstrated here aims to allow for the synthesis of a system that can successfully detect host pathogens in a matter of minutes. Furthermore, the proposition of modifying the internal capsid proteins (ICPs) of T7-like phage is presented for the first time in this work.

Several T7-like phage ICPs (proteins that structurally make up the interior of the phage capsid) are of particular interest here due their additional duty as ejection proteins (proteins that are ejected out of the phage upon host adsorption). Once the phage tail fibers adsorb to the host, the smallest ICP/ejection protein, gp6.7, is immediately propelled through the host membranes and into the cytoplasm, forming an initial pore before three larger ICP/ejection proteins, gp14, gp15 and gp16, partially unfold, pass through the phage portal and tail, and refold to form the ejectosome - a tunnel-like structure spanning the outer membrane, peptidoglycan layer and inner membrane [19]. The ejectosome subsequently assists with phage genome translocation from the capsid into the host cytoplasm in order for protein expression and phage propagation to commence [20]. For this work, gp6.7 and gp14 were selected as ICP fusion candidates due to the former being smallest ICP and the only one that is ejected into the host cytoplasm and the latter because it forms the outer part of the ejectosome and is therefore potentially available for pre-lysis exterior interactions with the extracellular solution.

The motivation for engineering the ICPs here is to develop a rapid diagnostic system for pathogenic bacteria by utilising the host-induced release of these proteins and the signal-generating enzyme fused to them. This hypothesised strategy holds immense promise for future diagnostic applications due to the reasons outlined below:

1. The enzyme (fused to the ICPs) is protected and silenced by the encapsulating barrier of the major and minor capsid proteins which form the icosahedral capsid [**?**], therefore no signal can be detected whilst the fusion remains internal.
2. The ICPs are expelled out of the phage upon host adsorption, thus the fused enzyme is also propelled out and exposed for signal generation, only when the host is present.
3. Previously published phage-based diagnostic models are entirely reliant on phage synthesis and accumulation inside the host and host-mediated transcription and translation of a reporter gene inserted into the phage genome - a prolonged processes. Here, the proposed model is based on pre-translated, structural fusion proteins that are encapsulated inside the phage capsid. This does not rely on intracellular host processing and only requires host presence, thereby enabling a truly rapid diagnostic procedure.

Following on from the conception of this strategy, the first key component to be considered is the choice of enzyme to fuse to the ICPs. The ideal candidate must be of appropriate size so that it can physically fit inside the phage capsid without having a significantly detrimental impact on the phage subsistence; it must not generate detectable signal whilst it resides within the capsid; it must have a certain degree of structural pliability so that it can be fused to an ICP without inhibiting its function; it must produce a specific signal that is not susceptible to background noise in a bacterial environment; and it must have significantly strong signalling capabilities so that even in small quantities it can be reliably detected.

When taking into account all of these considerations, the luciferase enzymes are presented as attractive options due to their high specificity and sensitivity. One particular luciferase-based bioluminescence platform that was commercialised in 2010, NanoLuc®, ticks many boxes and was chosen as the choice of fusion for this application. NanoLuc is a small (19.1 kDa) luciferase which catalyses the conversion of a novel coelenterazine analog (2-furanylmethyl-deoxy-coelenterazine or furimazine) into furimamide - a reaction that emits high intensity, glow-type luminescence [23]. This system has enhanced thermal and acidic stability, increased brightness and prolonged glow kinetics compared to other well-known luciferase candidates such as Firefly and Renilla [22]. The ability for NanoLuc to be detected at very low quantities (as low as 0.01 pM [24]) is also favourable. Furthermore, previous demonstrations that NanoLuc can be split into two spontaneously complementing sub-units that are not sensitive to unfolding/refolding cycles [25] (referred to here as C-terminal NanoLuc (CtNL) and N-terminal NanoLuc (NtNL)), which further decreases the size of the component to be packaged inside the capsid and compressed through the phage tail, cement its place as the ideal candidate for an ICP fusion.

### 3.2 Rationale

The initial aim of this project was to establish a rationale that could justify the further investigation of utilising *Podoviridae* family internal capsid proteins (ICPs) fusions as vessels for highly specific and rapid bacterial detection. The two ICPs focused on in this study are gp6.7 and gp14, and the enzyme chosen to be fused to the ICPs is an N-terminus sub-unit of NanoLuc (NtNL). In each phage particle there are 18 copies of gp6.7 and 10 copies of gp14 [19], therefore in an achievable phage stock titre of 10^10^ PFU/mL (10 billion phage particles per mL) it would be expected that there would be 180 billion copies of gp.6.7 and 100 billion copies of gp.14 present within the stock. In an engineered phage stock of the same titre, where either gp6.7 or gp14 is fused to NtNL at a modestly calculated success rate of 10% (1 billion successfully engineered phage particles per mL) and where a continuously conservative estimate of 10% for the number of engineered fusion protein copies per phage is applied (1.8 copies of gp6.7::NtNL or 1 copy of gp14::NtNL per phage), it can be estimated that there would be 1.8 billion copies of gp6.7::NtNL or 1 billion copies of gp14::NtNL present within the 10^10^ PFU/mL engineered phage stock.

Continuing with this rationale, the next calculation to consider is the minimum known number of NanoLuc copies required for detectable light emission. Furthermore, previously published data has shown that 0.01 pM of NanoLuc is capable of generating detectable light [24]. The molecular mass of NanoLuc is 19.1 kDa (19100 g/mol), which can be used to calculate its mass in a volume of 1 mL (0.001 L) at a molar concentration of 0.01 pM (10^*−*13^ mol/L):

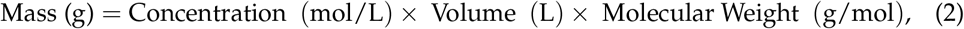

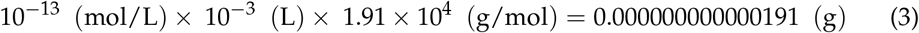

So, now that the minimum concentration of NanoLuc needed to emit detectable light (9.1 *×* 10^*−*14^ g/mL) is known, Avogadro’s constant (6.02214076 *×* 10^23^) can be used to calculate the number of NanoLuc copies in this concentration:

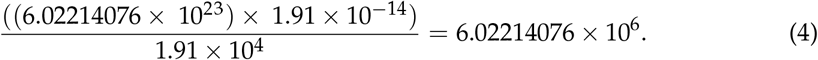

In conclusion, from these calculations we can deduce that 6,022,140 NanoLuc copies per mL is the minimum amount needed for the emission of detectable light. If the number of gp6.7::NtNL and gp14::NtNL copies per mL that were conservatively calculated above (1.8 billion and 1 billion copies respectively) are considered, and a final cautious estimation of 10% for the number of fusion proteins that successfully complement with their C-terminal (CtNL) counterpart is applied, this results in a final copy number of 180 million for gp6.7::NanoLuc or 100 million for gp14::NanoLuc per mL. Armed with these estimates, it can rationally be concluded that there would be a sufficient quantity of NanoLuc (>6 million copies) per mL in the phage-based pathogen detection solution.

### 3.3. Fusion design and testing

Once the rationale was established, work was initiated on designing, constructing and testing the fusion proteins. The simple design of the fusion comprises the phage ICP and NtNL joined by a flexible (Gly-Gly-Gly-Gly-Ser)2 linker. The N-terminus of NtNl is the end that is fused whereas the C-terminus of NtNL is exposed so that it can freely complement with the N-terminus of CtNL to form the full NanoLuc enzyme without obstruction. This fusion gene design was ordered and subsequently cut and inserted into the pBEST plasmid for protein expression (Figure 1). Using the Benchling ‘Analyze As Translation’ tool, it was calculated that the sizes of gp6.7::NtNL and gp14::NtNL were 15.5 kDa and 28.2 kDa respectively.

**Figure 1.**
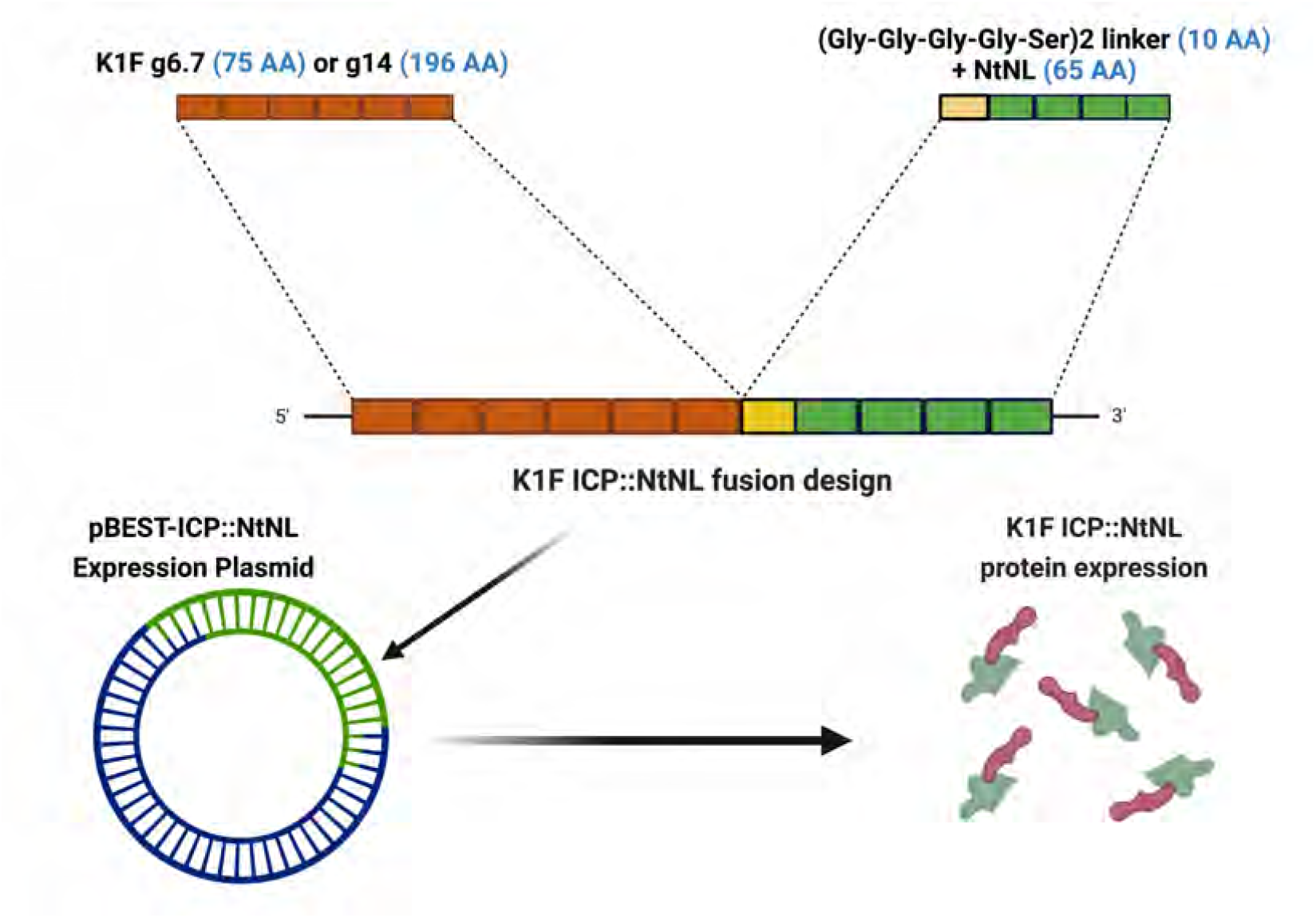
ICP::NtNL fusion protein design schema. The N-terminus of NanoLuc is fused to the C-terminus of the phage internal capsid protein with a (Gly-Gly-Gly-Gly-Ser)2 linker. This fusion sequence was inserted into the pBEST expression plasmid using restriction cloning, enabling expression of the fusion protein.

Once the fusion constructs had been successfully assembled into the expression plasmid, an in-house TXTL system that had been optimised for the pBEST plasmid was used to rapidly test the luminescence emission capabilities of the fusions. Also, by doing so, this would confirm that the act of fusing NtNL to the phage ICPs does not cause suppression of spontaneous CtNL complementation to form the full NanoLuc enzyme. The TXTL workflow displayed in Figure 2a allowed for rapid construct testing by simply expressing the various pBEST templates in TXTL reactions, followed by directly combining NtNL and CtNL reactions, and finally adding the furimazine substrate before measuring luminescence on the plate reader.

**Figure 2.**
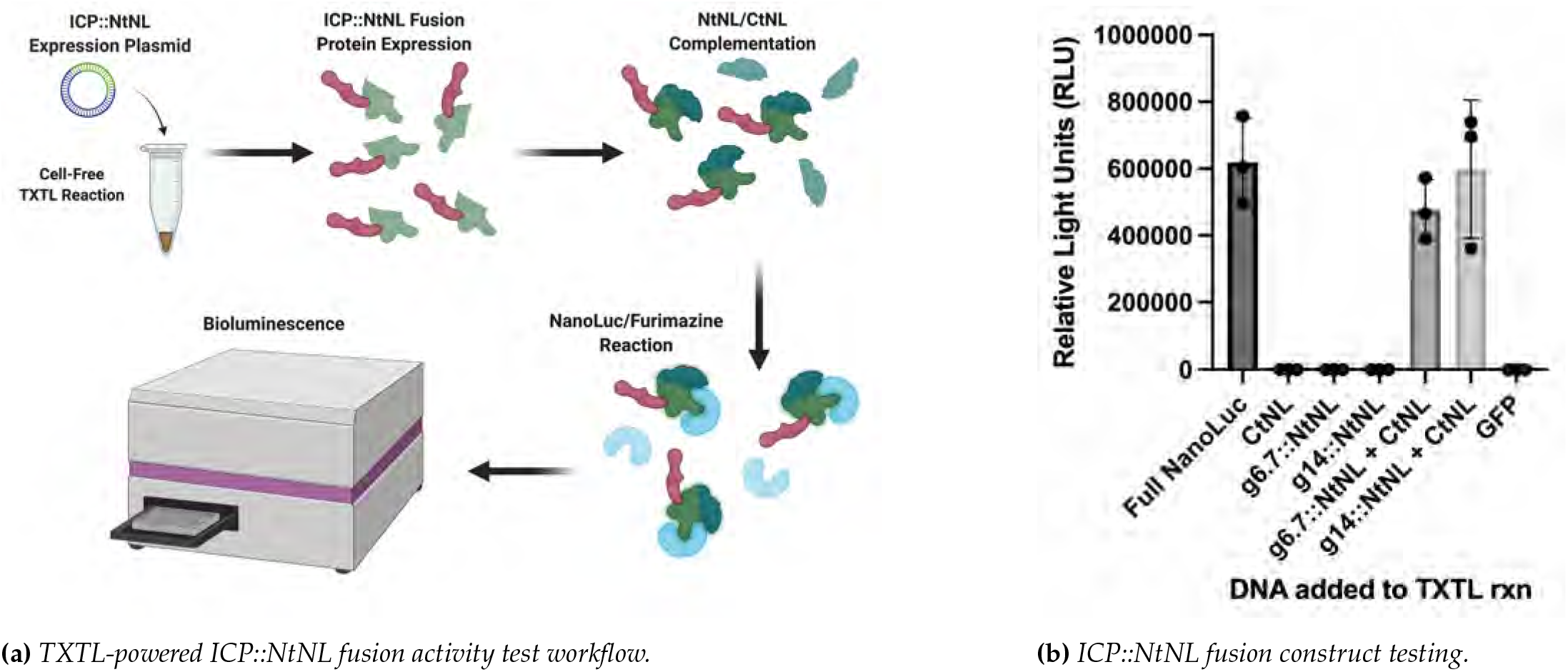
TXTL-powered ICP::NtNL fusion activity testing. **a:** Rapid testing workflow. The ICP::NtNL fusion protein was expressed in a TXTL reaction. The C-terminus of NanoLuc (also expressed in TXTL) was then provided to facilitate spontaneous NtNL::CtNL complementation. The NanoLuc substrate (furimazine) was added to the sample and luminescence was measured in a plate reader. **b:** ICP::NtNL fusion construct testing. Identical TXTL reactions were supplemented with 18mM of the pBEST plasmid inserted with: full NanoLuc, CtNL, g6.7::NtNL, g14::NtNL or GFP. The reactions were incubated for 8 hours and then immediately combined with an equal volume of Nano-Glo® assay buffer (containing furimazine). The fusion constructs were tested for luminescence both on their own and immediately after being combined with an equal volume of the CtNL TXTL reaction.

TXTL-testing the constructs confirmed that both ICP::NtNL fusions were capable of spontaneous complementation with CtNL and subsequently they both gave comparable luminescence measurements to the full NanoLuc control (Figure 2b). Following on from collecting this data and therefore confirming the fusion viability and activity, the next challenge - engineering the phage - could begin.

### 3.4 Cell-free TXTL synthesis of K1F phage

Prior to experimenting with the TXTL engineering method, it was first essential to successfully synthesize K1F phage in a TXTL reaction. As the demands of phage synthesis on a TXTL system are much higher than single protein expression, an optimal system is preferable. Therefore, it was decided to use the highly powerful myTXTL kit (Arbor Biosciences, Michigan, USA).

Figure 3 displays the data and images generated from optimising K1F synthesis in the myTXTL system. One early sign of a successful TXTL phage synthesis reaction (prior to validating via a spot test of plaque assay) is observing whether the reaction has become opaque or not. As shown in the upper tube in Figure 3c, the K1F synthesis reaction has caused the liquid to turn opaque, whereas the reaction in the bottom tube (a recombinant protein expression) has remained relatively clear. Further analysis via a plaque assay enables an accurate titer to be calculated. Optimal reaction conditions in myTXTL produced a very high K1F titer of approximately 10^12^ PFU/mL.

**Figure 3.**
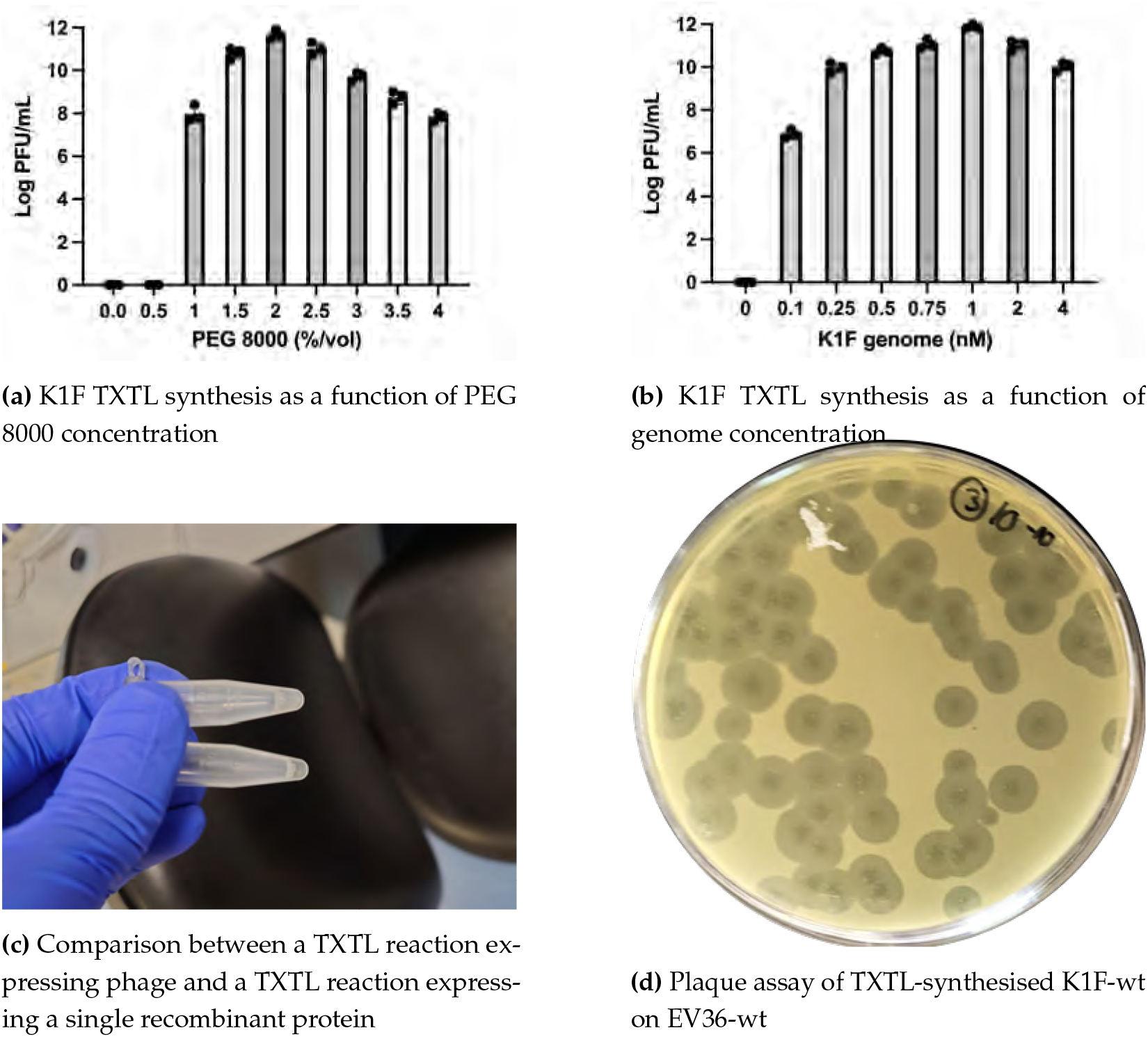
myTXTL optimisations for cell-free K1F-wt synthesis. The optimal PEG 8k %/volume (**a**) and K1F genome concentration (**b**) were found by testing a range of different values in the myTXTL system, (+/ SD, n = 3). The 12 *µ*L reactions were carried out in triplicate and incubated for 16 hours at 29 °C. Successful TXTL phage synthesis reactions typically appear more opaque (**c**). Plaque assays were carried out on a series of TXTL reaction dilutions with EV36-wt to calculate the phage titer (**d**).

The optimal amounts of PEG and K1F genomic DNA are also displayed. As Figure 3a shows, a minimum of 1%/vol of PEG 8k is necessary for the reaction to be viable, and when increasing PEG 8k from 1% - 2% (the optimal amount), a 10,000-fold increase in PFU/mL is exhibited - validating the importance of PEG 8k in cell-free reactions. Previous research has found the optimal genome concentration for T7 in a TXTL reaction to be 0.25 nM [27], however as shown in Figure 3b, the optimal genome concentration for K1F found here is 1 nM.

### 3.5 An investigative platform comprising cell-free TXTL and electron microscopy for studying phage synthesis

Following on from successfully optimising the myTXTL platform for K1F synthesis, a side project became of interest whereby it was hypothesised that the open and controllable cell-free environment would be well-suited for time-course imaging of phage. Traditionally in phage-related studies, electron microscopy (EM) would be used solely for capturing stand-alone images of phage in certain scenarios. However, this novel proposition enables the user to discover and track the characteristics of phage synthesis over the course of a controlled TXTL reaction whilst aligning the images with the corresponding phage titer for further insight. Furthermore, due to the highly controllable TXTL environment, it is possible to precisely quantify the start and end points of DNA expression (i.e. when the genome is added to the reaction and when the transcriptional inhibitor, rifampicin, is added to the reaction). This defined level of control is not easily attained *in vivo* due to the turbulent nature of phage propagation and opaque composition of membrane-encapsulated bacterial cells.

Figure 4 displays the proposed workflow, where cell-free TXTL phage synthesis reactions are stopped at various time intervals with the addition of 100 *µ*g/mL rifampicin - an RNA-polymerase deactivator which directly inhibits transcription and therefore any subsequent translation and phage synthesis processes within the reaction are also ceased. Then, 10 *µ*L of the inhibited TXTL reactions are directly applied to electron microscopy grids and imaged with EM - subsequently producing a ‘real-time’ image of each TXTL phage synthesis time interval reaction. By carrying out a plaque assay on each time interval reaction it is also possible to map the images to their corresponding phage titer.

**Figure 4.**
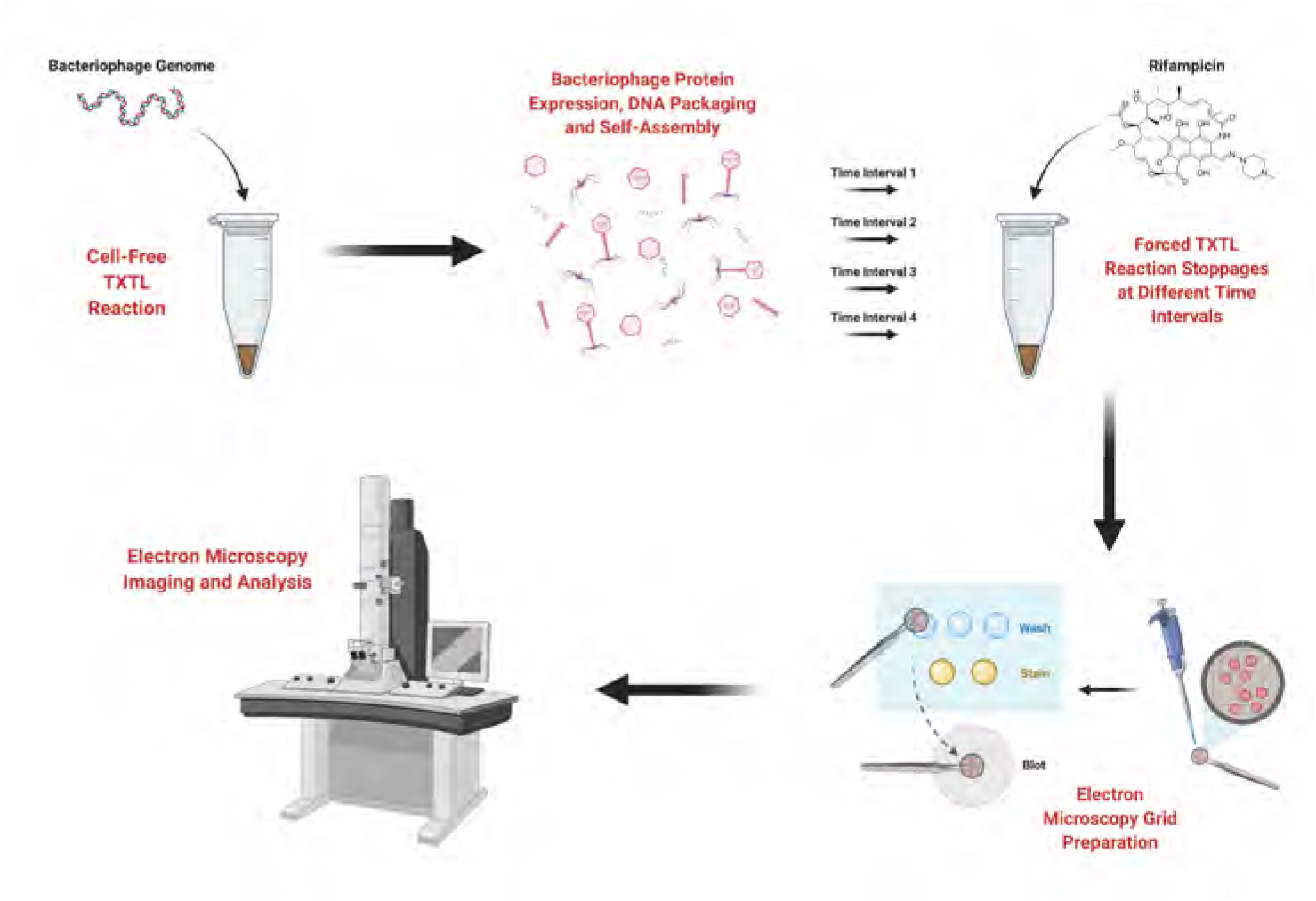
Workflow for TXTL synthesis-mediated electron microscopy phage assembly analysis. Phage are assembled in a cell-free reaction and the reactions are stopped at different time intervals. Each time interval reaction is then visualised via electron microscopy.

To display a representation of the different stages of TXTL phage synthesis (whilst considering the high cost of EM imaging), five time interval reactions were selected for imaging: 15 mins, 30 mins, 45 mins, 1 hour and 3 hours. As shown in Figure 5, the region of rapid K1F phage synthesis in TXTL spans the aforementioned time intervals and after 3 hours the phage titer begins to plateau between 10^11^ - 10^12^ PFU/mL. Between 0 - 1 hours a rapid increase in K1F synthesis is observed, with the PFU/mL rising from 0 - 10^7^. Then, from 1 - 3 hours a further 1000-fold increase in phage titer is observed. It was therefore of high interest to use this novel TXTL/EM approach to visually observe any changing characteristics in phage assembly or propensity over the course of these first 3 hours of TXTL synthesis.

**Figure 5.**
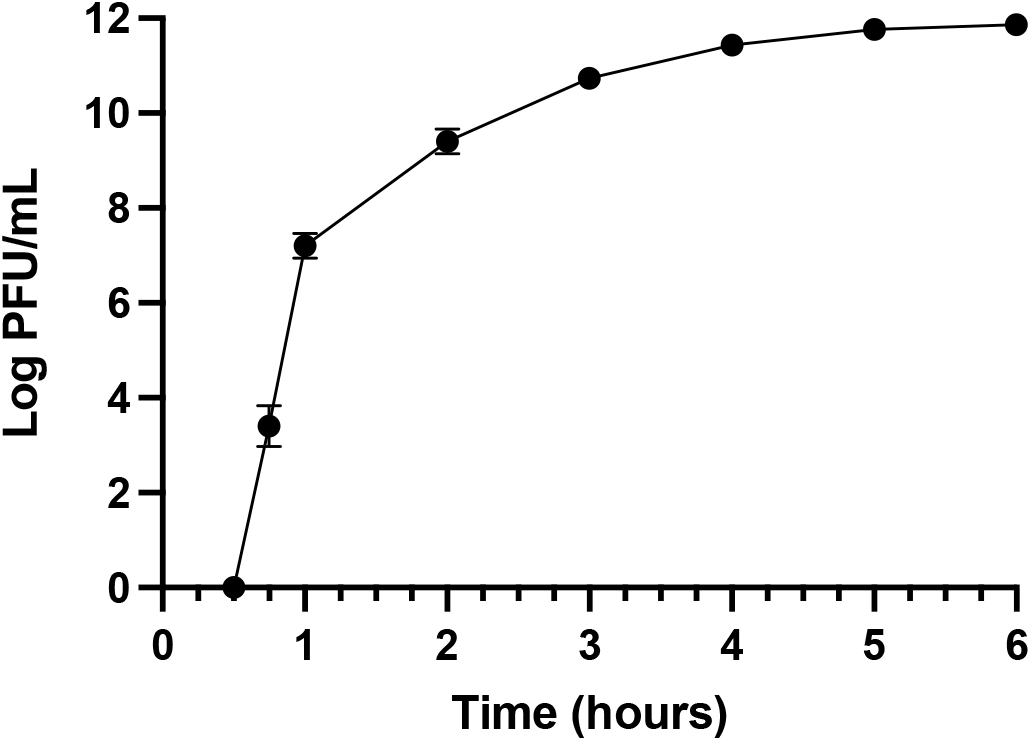
Kinetics of K1F TXTL synthesis. A concentration of 1 nM K1F genome and 2% PEG were added to the myTXTL master mix for this experiment. Plaque assays were carried out to calculate titers for each time interval. Each sample was done in triplicate. No phage is produced during the first 30 minutes of incubation. The first recorded PFU is at 45 minutes and K1F synthesis typically reaches plateau within 4-5 hours at approximately 1012 PFU/mL, (+/ SD, n = 3).

The first two K1F TXTL synthesis time interval reactions that were imaged were 15 and 30 minutes. No observations could be made 15 minutes after the reaction had started (Figure 6). The images captured mainly consisted of the grey/black ‘background noise’ and sporadic lipid/protein clusters that are typical of a standard *E. coli* lysate EM image. In fact, a small aliquot of an empty myTXTL reaction (i.e. not expressing any DNA) was also analysed via EM and this produced comparable images to those seen at the 15 minute time interval. This is not surprising though, given that no K1F TXTL synthesis PFU titer is observed until the 45 minute mark (Figure 5).

**Figure 6.**
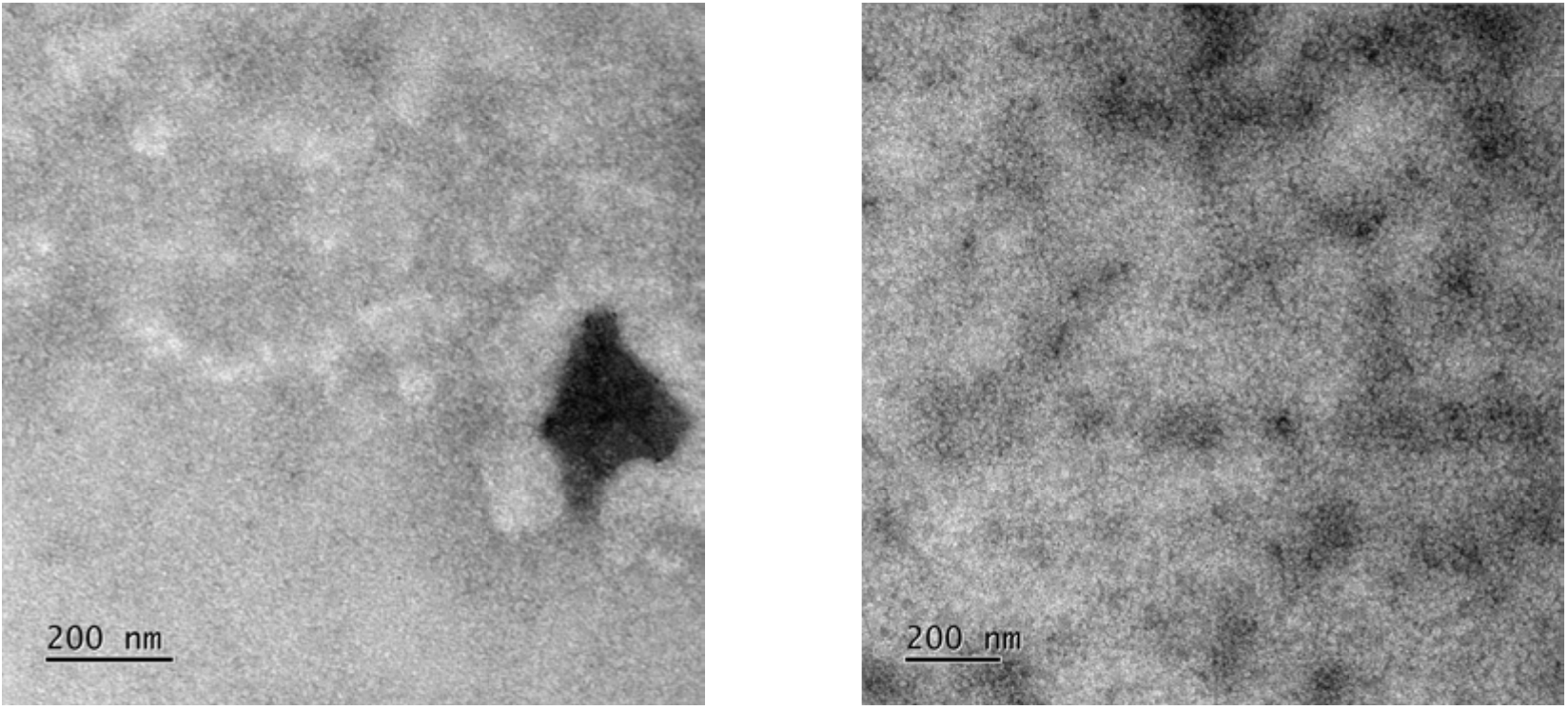
15 minute time interval: K1F TXTL synthesis visualised with EM.

A number of sporadic phage capsids could be seen at the half-hour mark (Figure 7), suggesting that protein expression and phage assembly were underway within 30 minutes of beginning the K1F TXTL synthesis reaction. The fact that still no PFU titer is observed at the 30 minute time interval (Figure 5) suggests that little or none of the phage capsids seen in these images represent viable phage, and will therefore be referred to as procapsids. As a typical component of dsDNA phage assembly and maturation, a procapsid is defined as a DNA-free phage capsid that is usually more spherical in shape and smaller than a matured capsid. Over the course of the assembly of a viable phage, the procapsid undergoes a dramatic conformational change whilst packaging the genomic DNA, which for phage T7, results in a larger, more angular capsid exhibiting the typical icosahedral structure [21] - this can be expected for phage K1F too. The identified procapsids can be seen within the black boxes in the two EM images displayed in Figure 7.

**Figure 7.**
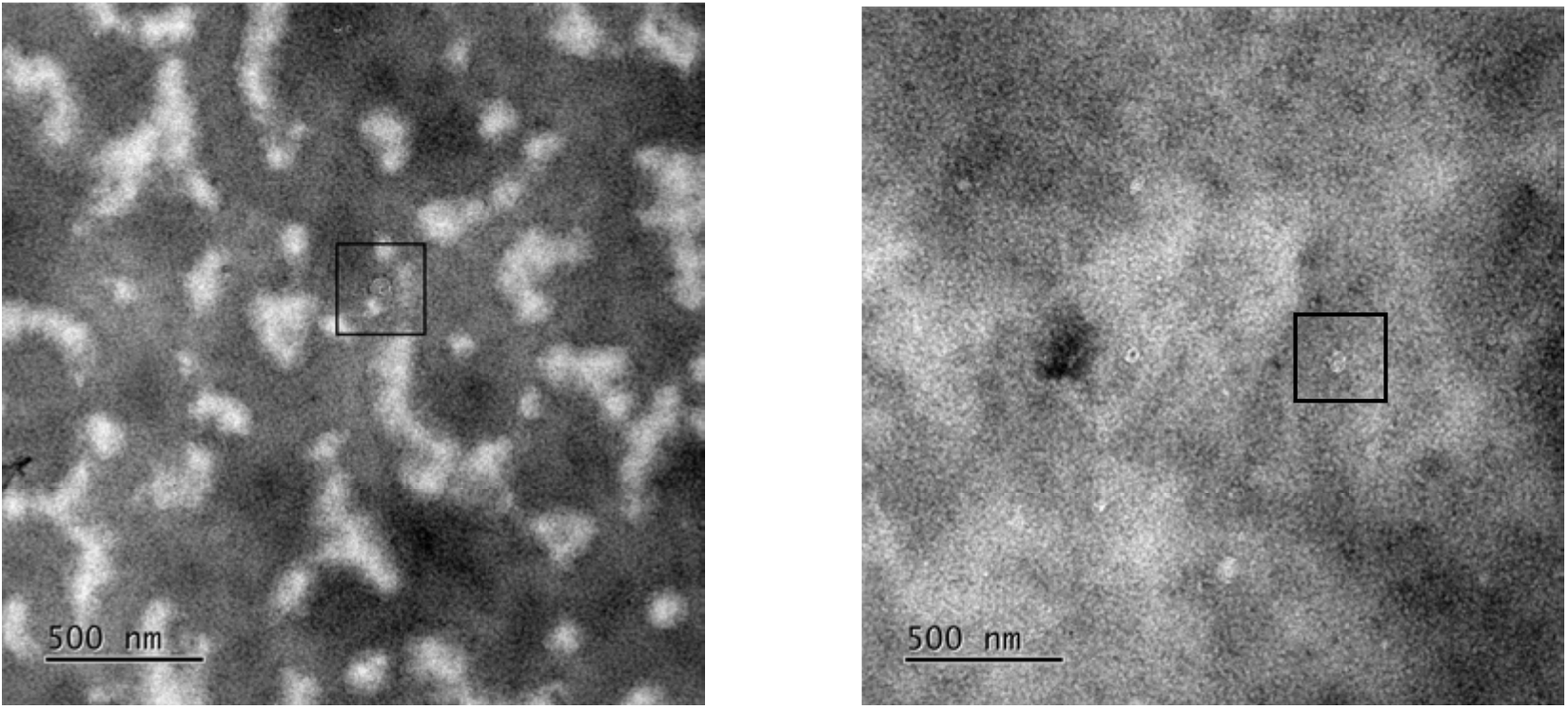
30 minute time interval: K1F TXTL synthesis visualised with EM.

Interestingly, for the 45 minute K1F TXTL synthesis time interval reaction - which corresponds with the first observable K1F phage titer of 10^3^ PFU/mL (Figure 5), EM analysis reveals that the procapsids appear to be accumulating together (Figure 8). One hypothesis for this procapsid accumulation phenomenon (PAP) is that, following on from phage protein expression and procapsid formation, the procapsids accumulate together whilst DNA packaging is undergone. Perhaps the DNA concatemers that are typical of T7-like phage serve as the rendezvous point for the PAP. Again, the events of interest are highlighted by the black boxes in the images shown in Figure 8.

**Figure 8.**
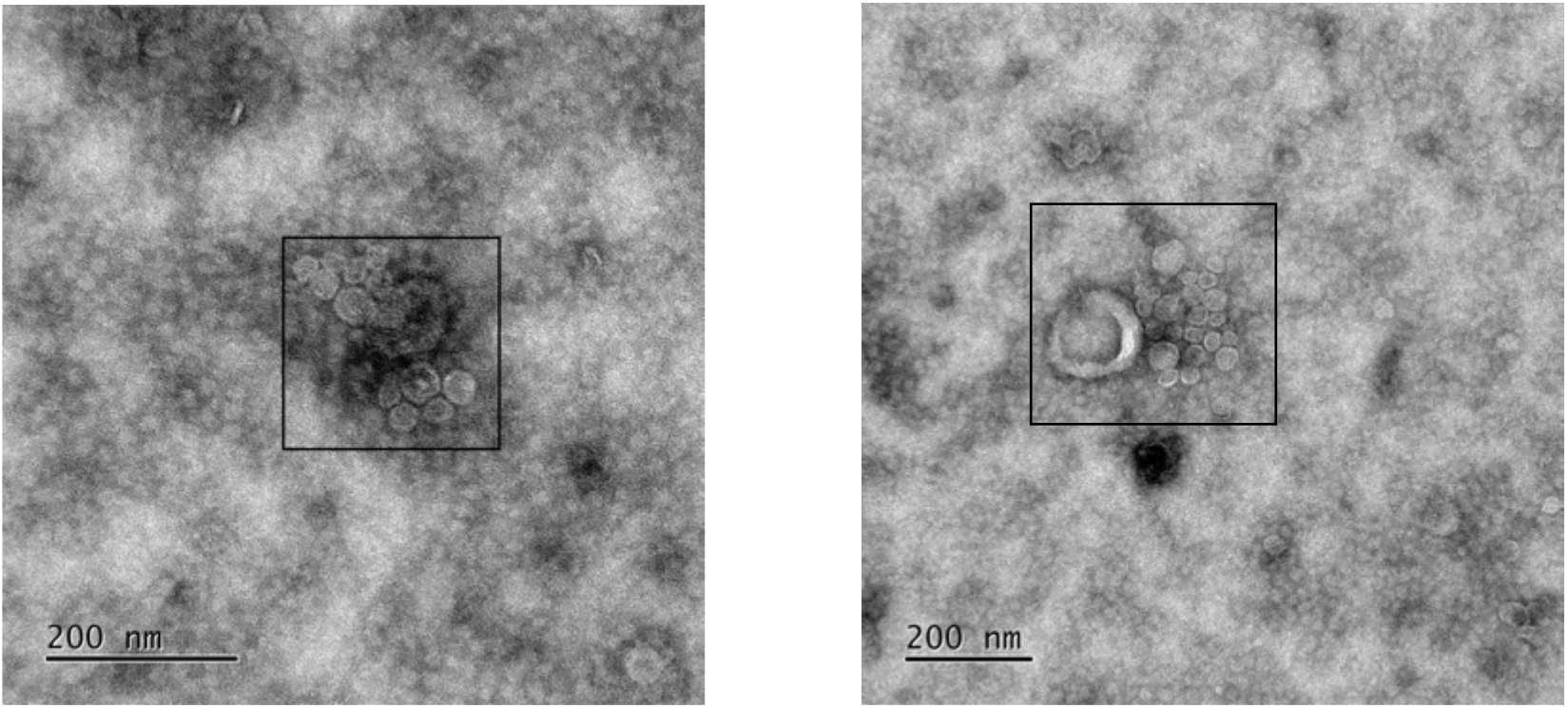
45 minute time interval: K1F TXTL synthesis visualised with EM.

The 1 hour time interval displayed in Figure 9 exhibits a continuation of the proposed PAP. Procapsid accumulation was observed at an increased abundance during EM imaging of these samples compared to the 45 minute samples. Furthermore, the 1 hour K1F TXTL synthesis time interval reaction aligns to a titer of 10^7^ PFU/mL (Figure 5). One interpretation of this data could suggest that the events displayed in the 45 and 60 minute images are in fact ‘DNA packaging’ events. This would suggest that premature phage accumulate together in large DNA packaging events as the last stage of their development and after which, they become viable phage. Furthermore, the increased abundancy of the PAP, and therefore possible DNA packaging events, from the 45 minute - 1 hour time interval could represent an acceleration of capsid maturation and subsequently viable phage synthesis. Indeed, the result of procapsid DNA packaging is in fact the generation of viable phage [21]. The rapid shift from 0 PFU/mL at 30 minutes, to 10^3^ PFU/mL at 45 minutes, to 10^7^ PFU/mL at 60 minutes would support this conclusion.

**Figure 9.**
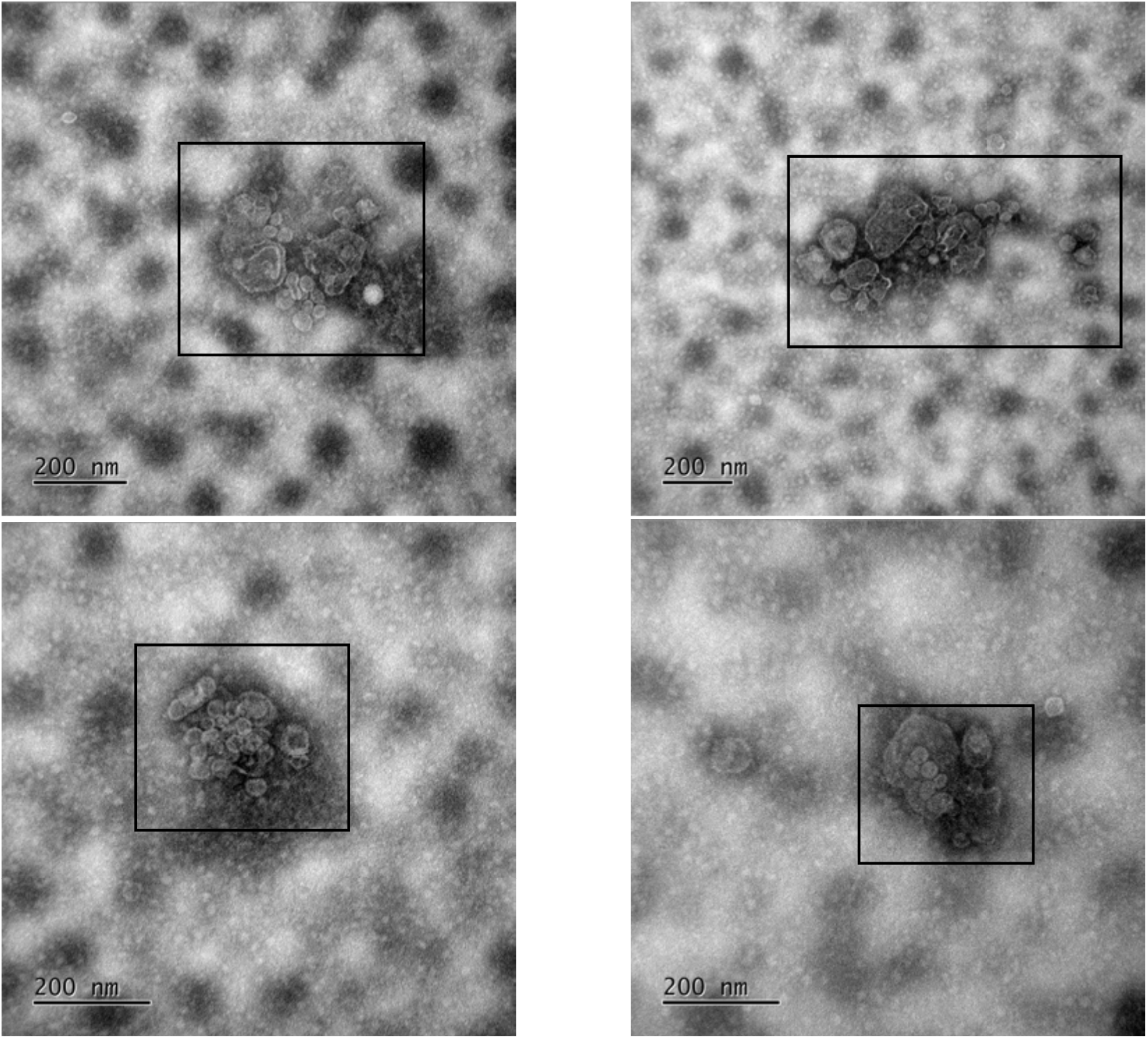
1 hour time interval: K1F TXTL synthesis visualised with EM.

The EM imaging results for the final K1F TXTL synthesis time interval reaction - 3 hours - display an entirely new characteristic not seen during imaging of the previous time interval reactions. The abundancy of phage particles can be seen to have increased, however, they appear to be much less accumulative (Figure 10). This suggests that the PAP and DNA packaging events are mostly complete by the 3 hour mark and that almost all phage synthesised in the reaction are viable at this time interval - this interpretation is supported by the fact that the titer observed here is 10^10^ PFU/mL and over the course of the following 5 hours phage synthesis decelerates and plateaus at approximately 10^12^ (Figure 5).

**Figure 10.**
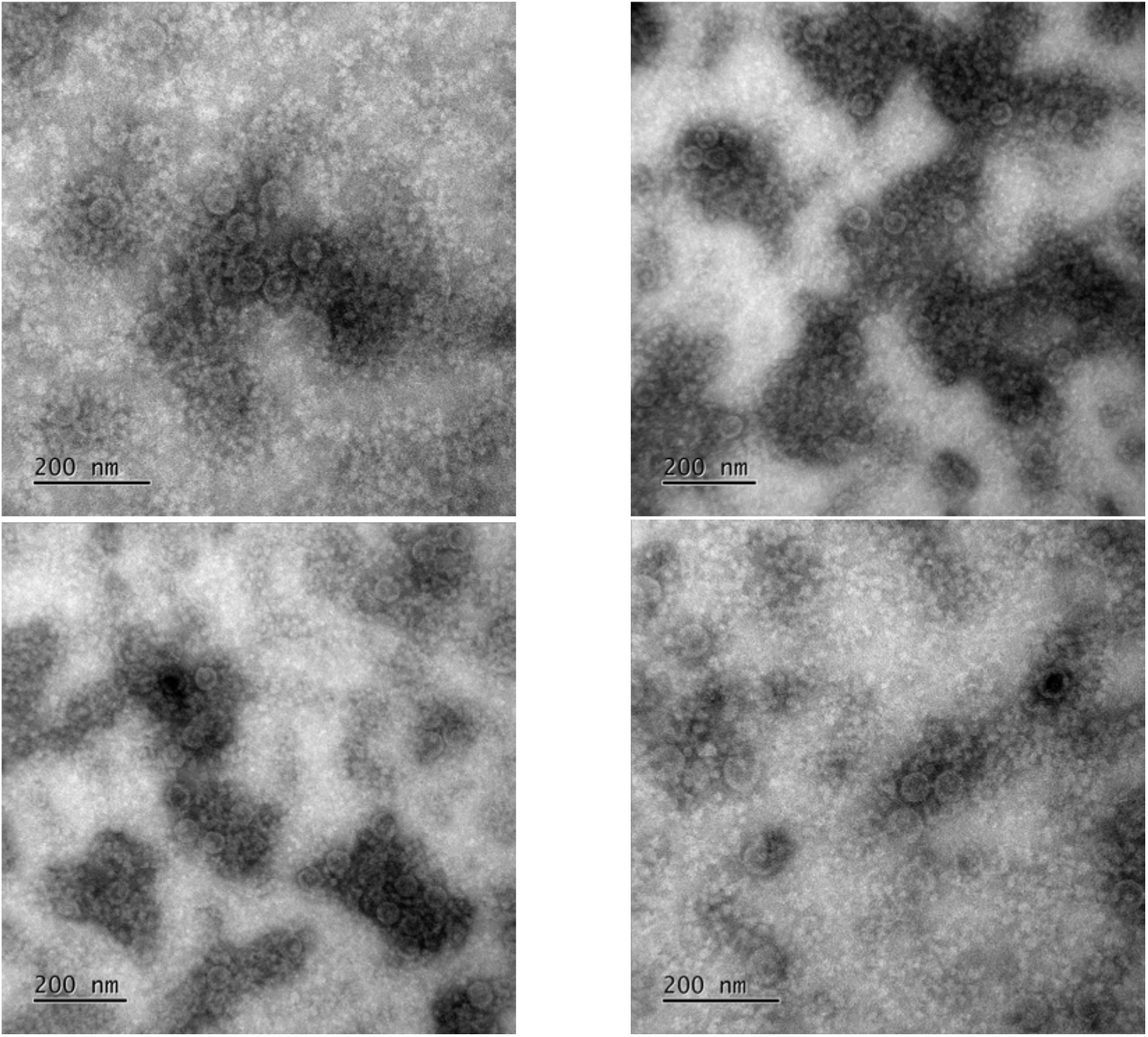
3 hour time interval: K1F TXTL synthesis visualised with EM.

Moreover, the phage displayed in the images in Figure 10 appear slightly larger (approximately 50 nm in diameter) than at previous time intervals and also exhibit a more uniform and angular conformation, which is typical and would be expected of a mature K1F capsid. The lack of the PAP in these images is not surprising as, at this plateauing stage of K1F phage synthesis within the TXTL reaction, the majority of the phage have already been synthesised and matured and the transcriptional and translational machinery alongside the energy reagents within the cell-free reaction have been exhausted almost to completion. Therefore, it is not expected that a significant amount of new procapsid formation and accumulation would be observable compared to the amount of viable phage already present.

A deep search of the literature revealed that, whilst various mentions of procapsid accumulation are found in an array of decades worth of research papers, they all seem to refer to a process whereby defective capsids which are unable to package their DNA accumulate together. This suggests that the PAP is in fact a known process (albeit, perhaps unsurprisingly, it hasn’t previously been referred to specifically as “the PAP”), however, it is not thought to be responsible for that which was hypothesised in the previous paragraph (i.e. DNA packaging) - seemingly quite the opposite. For example, in a publication from the mid 1970s [28], it is described and visually demonstrated that the procapsids of P22, a phage that infects *S. typhimurium* which is actually very similar to T7 and K1F, accumulate amid “vegetative DNA” concatemers in a way that is comparable to that which is shown in Figures 8 and 9. More recently, it has also been shown that when SPP1, a phage that infects *B. subtilis*, is engineered so that it cannot package DNA into its capsid, the procapsids which are generated accumulate together [29] and for defective *ϕ*12, a *S. aureus*-infecting phage, where the disarming of DNA packaging machinery, again, leads to procapsid accumulation [30].

However, if the PAP events are an exclusive occurrence for defective phage, then it is surprising that none of these events could be found at the 3-hour mark (Figure 10), as there is no reason why they would disappear. One explanation for what is occurring, now hypothesised after considering the results found in the literature alongside the EM/TXTL findings, is that the PAP occurs universally for all phage and it is in fact representative of DNA packaging events (regardless of whether those packaging events are successful or not). The reason why it has previously been described as a characteristic of defective phage [28– 30], is because these phage are not capable of progressing past the DNA packaging stage of maturation and therefore remain in the accumulative state. However, it is hypothesised here that “normal” phage also go through the PAP phase whilst they gather around DNA concatemers in a collective (perhaps cooperative? e.g. sharing packaging machinery and/or ATP energy) DNA packaging event. Furthermore, once they have successfully packaged their DNA and become mature phage there is no longer a necessity for accumulation and they progress to the mature phase captured in Figure 10. This concludes the K1F TXTL synthesis investigative detour and the remaining results will revert back to the ICP phage engineering investigation.

### 3.6 in vitro non-genomic TXTL ICP phage engineering to generate single-use, host-dependent diagnostic phage

A fast and simple non-genomic phage engineering strategy was devised whereby cell-free TXTL is utilised for the expression of the ICP::NtNL fusion *and* phage genome simultaneously, which can conceivably generate a packaged diagnostic phage. This approach is preferable compared to an *in vivo* approach due to the refined and controllable environment of cell-free alongside its superior protein expression capabilities. This approach is visualised in Figure 11. As shown in this figure, there are two possible TXTL phage engineering approaches - the first comprising the simultaneous expression of the phage genome and pBEST-ICP::NtNL plasmid and the second expressing the phage genome whilst supplying pre-TXTL’d ICP::NtNL fusion protein as a reaction additive. It was planned for both methods to be attempted in order to find the optimal approach, whilst considering both the packaging efficiency and phage titer generated. The two approaches are referred to as: endoTXTL (endogenous and simultaneous expression of phage and fusion DNA) and exoTXTL (phage DNA expression supplemented with exogenous fusion protein). Plaque assay results showed that the endoTXTL engineering attempts successfully yielded viable phage, however, exoTXTL engineering attempts failed to generate any phage and all subsequent repeats using the exoTXTL method were also unsuccessful (data not shown).

**Figure 11.**
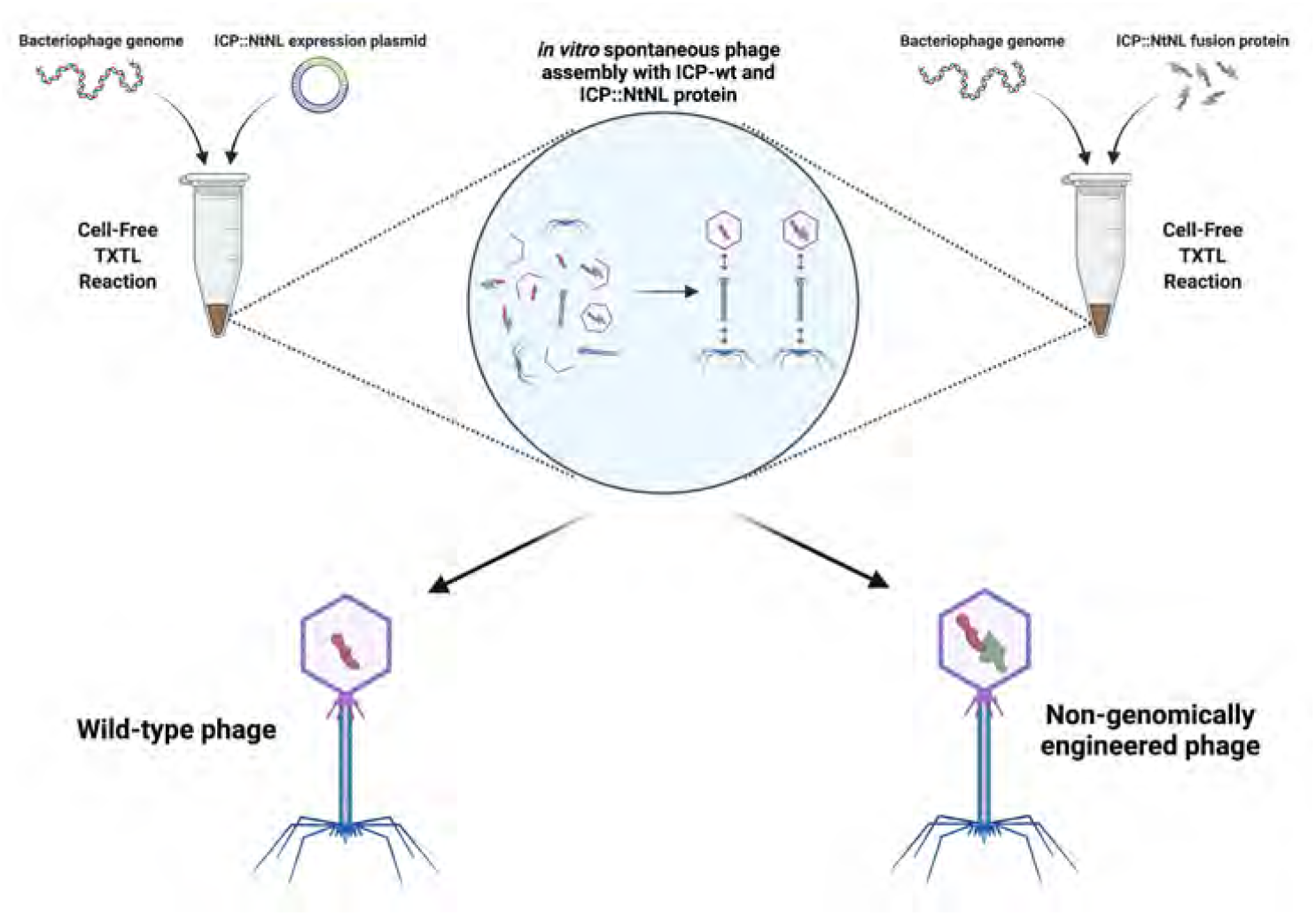
Non-genomic TXTL ICP phage engineering strategy used for packaging the NtNL fusion protein into the phage capsid. Phage are packaged in a cell-free reaction either with co-expressed fusion protein (endoTXTL) or pre-expressed fusion protein (exoTXTL), generating a mixed population of WT and non-genomically engineered phage.

A possible explanation for this is that the presence of the fusion protein at the start of the exoTXTL reaction may have interfered with the molecular crowding equilibrium maintained by PEG 8k or otherwise destabilised the TXTL ecosystem leading to the failure of K1F synthesis. The PEG 8K-mediated molecular crowding within TXTL is especially important for phage synthesis [27] and so any interference with this is likely to be detrimental for the reaction. Subsequently, rather than spending a significant amount of time on attempting to extensively optimise the exoTXTL approach (e.g. by adding different exogenous protein concentrations and/or purifying the protein before adding it and/or adjusting the PEG 8k concentration), it was decided to proceed with the endoTXTL phage engineering approach.

The endoTXTL approach was used to generate the following engineered phage: TXTL-K1F*gp6.7::NtNL* and TXTL-K1F*gp14::NtNL*. Aliquots from several stages of the phage engineering process were taken and tested for luminescence with the Nano-Glo® assay (Figure 12). Hypothetically, this method could be used to manufacture single-use, host-dependent diagnostic phage (or any other engineered phage generated using the method) at industrial scale. The only limiting factors are, firstly, the size and cost of TXTL reaction that the user is capable of processing, and the secondly, the size of the filtration unit they can source, however, if a bespoke solution was built then this method could be used to produce a vast amount of single use phage.

**Figure 12.**
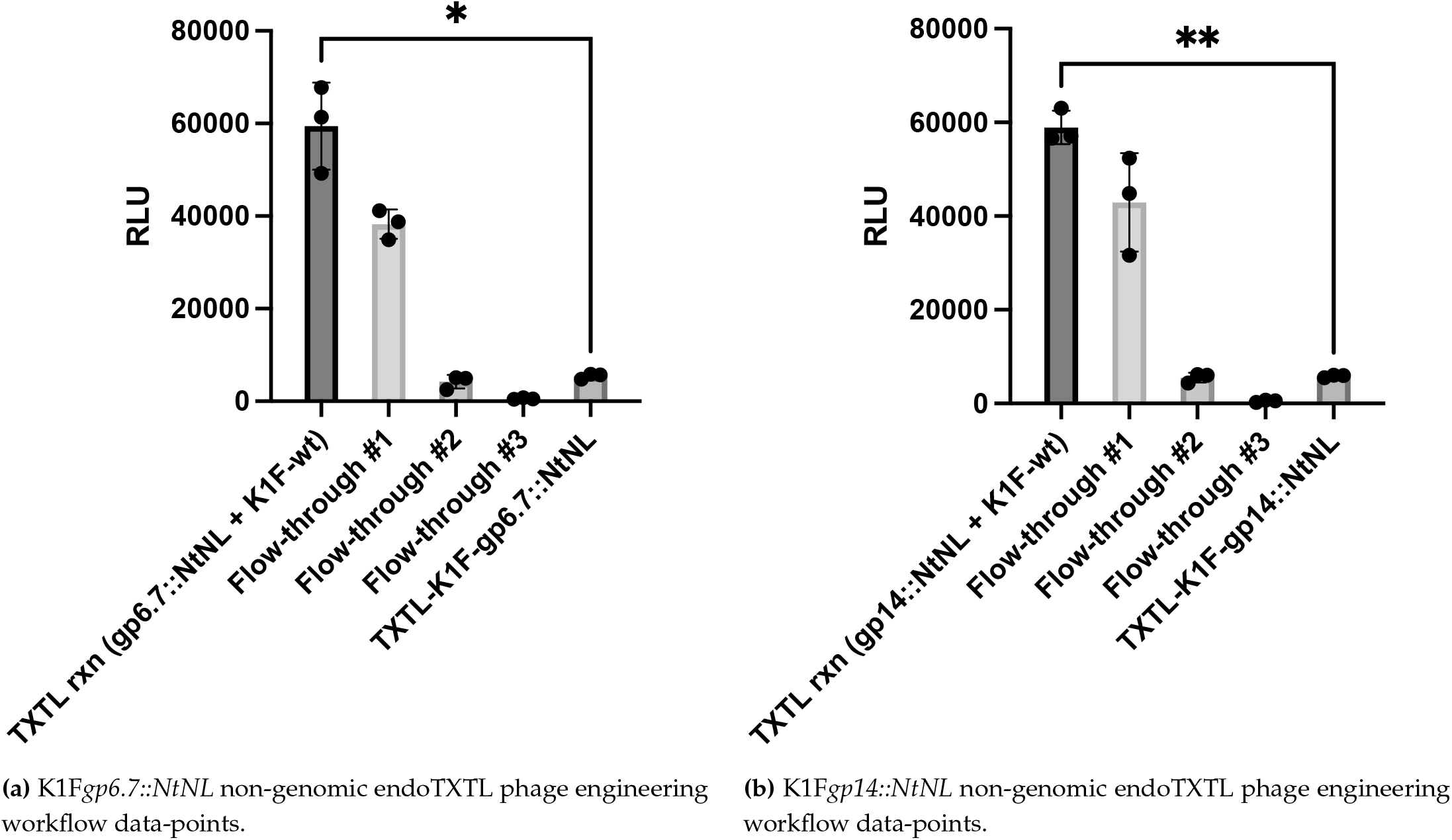
Non-genomic TXTL engineering workflow data-points. Luminescence was measured on aliquots taken at several stages during the **a:** K1F*gp6.7::NtNL* and **b:** K1F*gp14::NtNL* non-genomic endoTXTL engineering process. Flow-through samples were taken after passing the phage through a 100 kDa Amicon® Ultra filtration device three times, using a fresh device each time. All samples were combined with an equal volume of a CtNL TXTL reaction and left to incubate for 1 hour at room temperature to allow for spontaneous CtNL::NtNL complementation prior to the Nano-Glo® assay, (+/ SD, n = 3).

The data shown in Figure 12a and Figure 12b are largely comparable to one another. Data-points 3, 4 and 5 are as expected, with the first filtration flow-through maintaining the majority of the signal and the next two emitting only baseline luminescence. However, contrary to the logic that the 15.5 kDa (gp6.7::NtNL) and 28.2 kDa (gp14::NtNL) fusion proteins should entirely pass through the 100 kDa filtration unit, it would appear that a portion of the protein was in fact retained alongside the phage. Furthermore, the sixth data-point exhibits a slight increase in luminescence above the baseline level observed in the second and third flow-through samples. It was therefore essential that this background signal was taken into account during the subsequent testing of the diagnostic phage.

### 3.7 Heat-induced signal release by in vitro non-genomically TXTL engineered phage

After carefully sifting through the literature and finding relevant data concerning the expulsion of T7/K1F genomic DNA and other internal capsid contents, a quick acid test to verify successful signal packaging was devised. This was based on the findings that, upon heating to temperatures between 65°C - 80°C, it has recently been shown that phage T7 prematurely ejects its genomic DNA [31]. As the ejection of the ICPs precede DNA ejection [32], it is also hypothesised here that the phage ICPs are prematurely expelled at high temperatures too. Therefore, if successful ICP::NtNL fusion packaging had taken place, an increased luminescent signal should be detected at high temperatures when the internal capsid contents (containing ICP::NtNL) are heat-ejected out of the phage. However, before testing this, it was first important to observe what impact heating to different temperatures had on the ICP::NtNL fusions’ ability to successfully complement with CtNL and emit light when provided with substrate.

The data in Figure 13 reveal that, at higher temperatures (45 °C - 90 °C), the activity of ICP::NtNL is significantly diminished. Furthermore, the luminescent readout gradually decreases as the temperature is increased from 45 °C to 90 °C. This could be attributed to the high temperature-induced denaturation causing a decrease in the fusions’ ability to either 1). complement with CtNL or 2). catalyse the furimazine to furimamide reaction once it has complimented with CtNL to form the full NanoLuc enzyme, or 3). a combination of the two. The RLU values are similar at RT and 30 °C, suggesting negligible protein denaturation had occurred, which is to be expected. Data for gp6.7::NtNL and gp14::NtNL are comparable. Interestingly, previously published data suggest that the melting temperature of the full NanoLuc enzyme is 58 °C [24]. Moreover, the data presented in Figure 13 would suggest that, when the enzyme is split into two sub-units, its melting temperature is reduced to approximately 45 °C. This is suggested by the drastic reduction in RLU output from 30 °C to 45 °C, eluding to the enzyme activity and thus, its composition. It, perhaps, is not surprising though that the thermostability of NanoLuc is reduced when split into sub-units, as the optimal stability is to be expected when the fully formed and folded protein composition is displayed.

**Figure 13.**
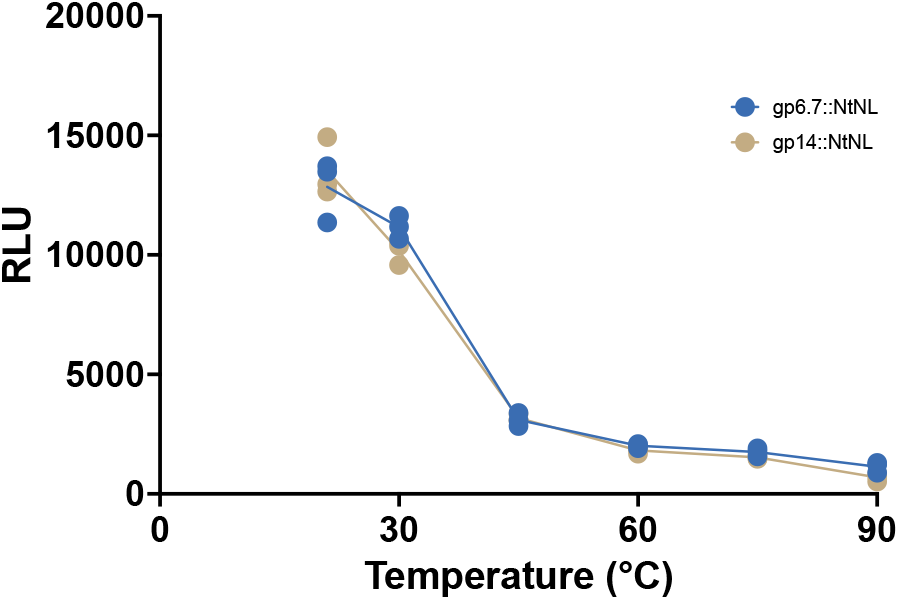
ICP::NtNL fusion thermal stability. TXTL-expressed fusion proteins (gp6.7::NtNL - blue or gp14::NtNL - gold) were exposed to different temperatures (RT, 30 °C, 45 °C, 60 °C, 75 °C and 90 °C) on a heat block, then let to return to RT. The samples were then combined with an equal volume of TXTL-expressed CtNL and let to incubate for 1 hour before carrying out the Nano-Glo® assay to measure their luminescent activity.

The ICP::NtNL thermal stability data from Figure 13 must be considered when analysing the heat-induced ejection results displayed in Figure 14. If the non-genomic engineering experiments had failed and the only ICP::NtNL fusion content present in each sample was unpackaged background (i.e. external to the phage capsids) then it would be expected that at higher temperatures (45 °C - 90 °C), the signal generated would be lower than at 30 °C and RT, due to the high temperatures denaturing the fusion protein (as displayed in Figure 13). However, the results in both Figures 14a and 14b show that when the engineered phage were heated to 75 °C, an increased signal compared to the RT sample is observed The room temperature (RT) values in Figures 14a and 14b represent the background luminescence present in the external solution surrounding the phage particles for TXTL-K1F*gp6.7::NtNL* and TXTL-K1F*gp14::NtNL* respectively. When exposed to 75 °C heat, both phage exhibited a statistically significant rise in RLU output compared to the RT values - indicating the confirmation of heat-induced release of encapsulated ICP::NtNL fusion from the phage and therefore suggesting the endoTXTL engineering experiments were a success.

**Figure 14.**
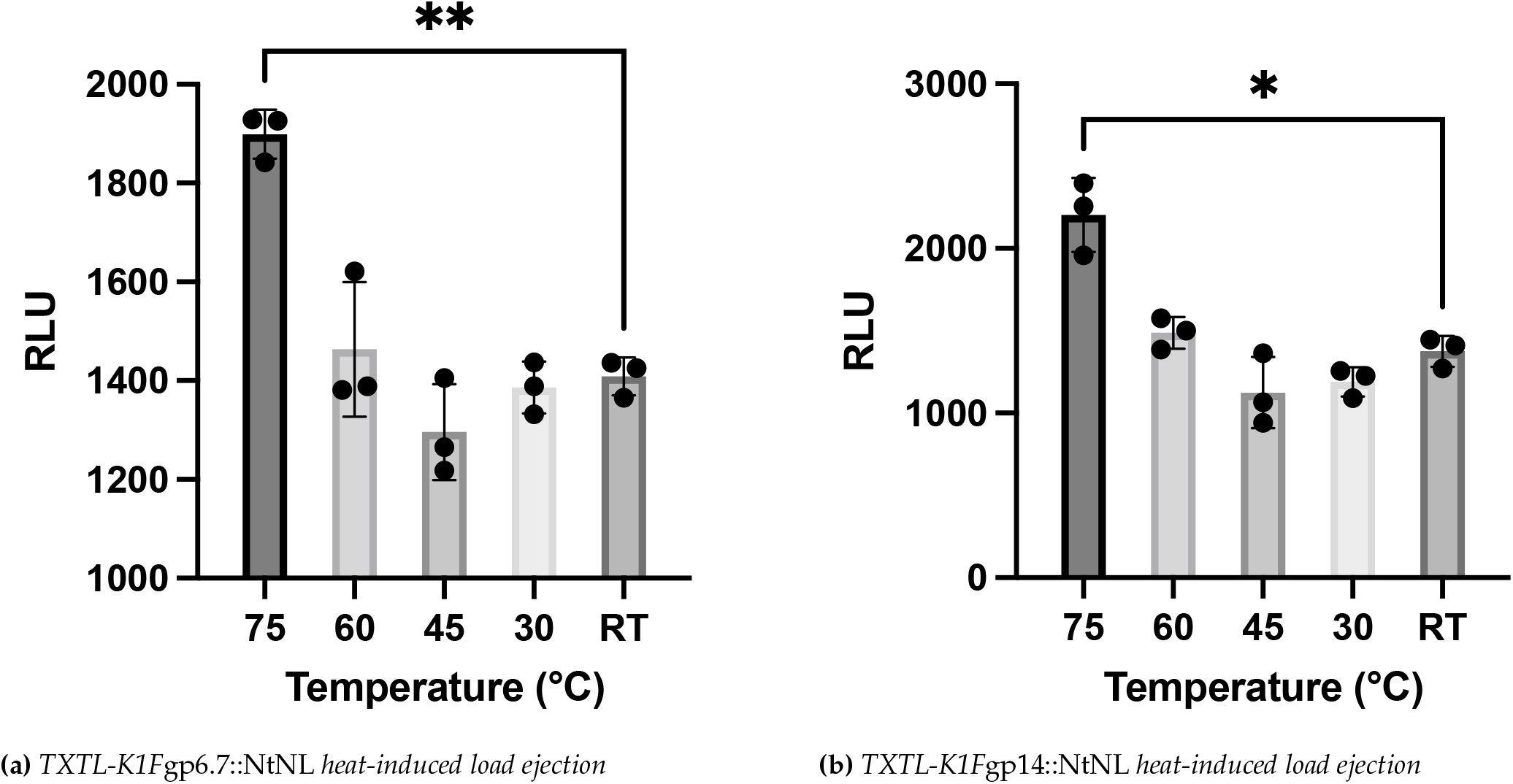
Luminescence measurements after heating non-genomically TXTL engineered phage to different temperatures. All samples were exposed to the desired temperature, then allowed to return to room temperature, then combined with an equal volume of CtNL TXTL reaction and left to incubate for 1 hour at room temperature to allow for spontaneous CtNL::NtNL complementation prior to the Nano-Glo® assay, (+/ SD, n = 3).

For both phage, when heated to 30 °C the RLU output decreased slightly but was comparable to the RT value. Moreover, heating to 45 °C incurred a further observable decrease in signal - suggesting the rising temperature was having a denaturing effect on the ICP::NtNL fusion. When the phage were subjected to an increased temperature of 60 °C, however, the mean RLU values increased to slightly above the background RT level, when it would be expected that the signal would continue to decrease as the temperature increases (Figure 13). This strongly suggests that a significant portion of fusion protein that was previously undetectable (i.e. packaged inside the capsid) had suddenly been released and become available for CtNL complementation, therefore increasing the signal output despite the enhanced denaturing capacity facilitated by the higher temperature. The considerable increase in RLU at the 75 °C temperature, where NtNL should be significantly denatured, adds further confirmation that the fusion had been packaged.

Next, to further consolidate the acceptance that the fusion proteins had been successful packaged, a SYBR green assay (Thermo Fisher Scientific, Waltham, USA) was carried out to confirm the release of DNA (indicating the simultaneous release of ICPs) at high temperatures. SYBR green is a dsDNA binding dye which, when bound to dsDNA, emits fluorescence. Therefore, the more dsDNA present, the higher the fluorescent signal. The results generated from this assay are displayed in Figure 15.

**Figure 15.**
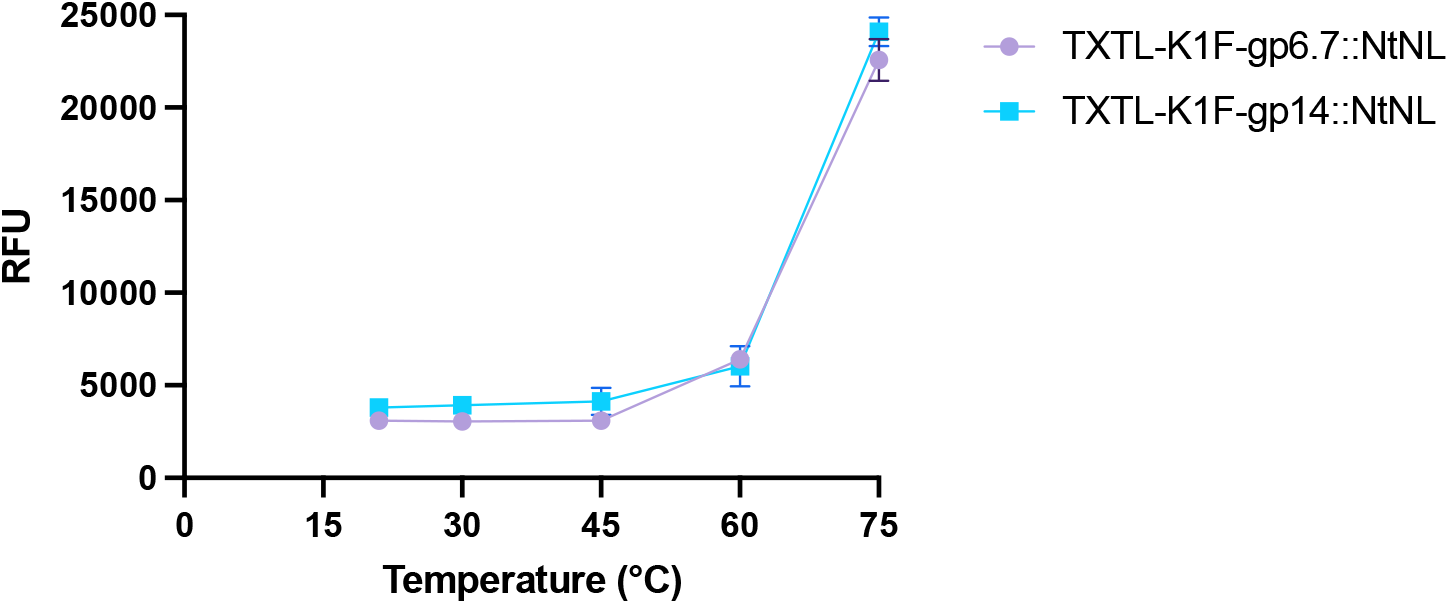
Fluorescence measurements after heating non-genomically TXTL engineered phage to different temperatures. All samples were exposed to the desired temperature, then allowed to return to room temperature, then mixed with the SYBR green reagent before being measured for fluorescence on a plate reader, (+/ SD, n = 3).

At temperatures ranging from RT to 45 °C, a small consistent RFU value is emitted, which can be considered as the background value. However, when the phage were heated to 60 °C, the fluorescence measurement slightly increased above the background value, potentially indicating that a small amount of DNA had been released from the phage. It is possible that this small amount of DNA represents the preliminary nucleotides which initially get ejected upon host adsorption. Usually, the rest of the genome is “pulled” into the host cytoplasm by the cytoplasmic RNA polymerases [33], however, in this case (i.e. in Figure 15) there are no RNA polymerases present as the ejection is heat-induced rather than host-induced. Therefore, it is reasonable to suggest that the SYBR green units are only able to bind to the small amount of preliminarily ejected DNA and subsequently, only a small amount of fluorescence is emitted. Alternatively, this slight increase in RFU could also represent a very small proportion of the phage prematurely ejecting their entire genome. At 75 °C, the RFU value significantly increases and clearly demonstrates the full release of the genomic DNA from a large proportion of the phage population. This was to be expected, as it has previously been shown for phage T7 [31].

Relating back to the heat-induced fusion protein ejection data shown in Figure 14, it can be hypothesised that, at 60 °C, a small proportion of phage are compelled to release their DNA, and therefore their ICPs and NtNL::ICP fusions would precede this. Furthermore, at 75 °C, a large proportion of the phage population are forced to eject their genomic by the high heat pressure and subsequently, a significant amount of the encapsulated fusion is released.

Following on from gaining these initially promising results, it was now possible to approach the host-induced signal release experiments knowing that, at least, it was highly likely that the fusion proteins had been successfully packaged. These experiments will be carried out and reported on in future work.

## 4. Discussion

The key results displayed in this work present a novel phage ICP fusion construct (Figure 1), a simple, TXTL-powered method for engineering phage (Figure 11), and a proof-of-concept diagnostic model where engineered phage emit signal in an inducible manner which mimmics host-mediated ICP ejection (Figure 14). The amalgamation of these results, alongside a literature review, suggest that the ICP::NtNL fusion is successfully packaged during the TXTL-powered non-genomic phage engineering process and subsequently retained within the capsid of the phage until induced, heat-mediated ejection (which mimics host-mediated ejection) occurs - an event which initiates the ejection of the ICPs, genomic DNA and any encapsulated ICP::NtNL fusion protein. The novel EM/TXTL phage assembly analysis workflow also presented in this publication offers a new possibility for phage investigation whereby researchers can observe the step-by-step progression of synthesis in a highly controllable environment without disturbing the sample. Future applications could allow for the manipulation of a plethora of parameters (only limited by the researchers imagination) to be visualised in real time and potentially, the uncovering of unexpected findings (such as the hypothesised PAP-mediated DNA packaging events).

With regards to the impact of the key results presented in this publication, future experiments will determine whether these heat-mediated ICP ejection results are reproducible in a host-mediated ejection setting. These experiments will represent a first step towards building a true *in vivo* test-at-home detection device. Furthermore, it is imperative that these further experiments are conducted to display a diagnostic model where CtNL can be provided extracellularly along with the other test components (i.e. lysis buffer and NanoLuc substrate - furimazine). One concern with this proposition is that the ejected fusion proteins will be degraded by endogenous host proteases before it is possible to expose them to the CtNL and furimazine. To counteract this, a pragmatic approach would be to lyse the cells at the earliest possible time-point with a lysis buffer incorporating a protease inhibitor cocktail in an attempt to free the ejected fusion into a protected environment before the entirety of it is degraded. It has previously been shown that the T7 phage eclipse time (i.e. the minimum time required for the host to produce the first phage progeny post-infection) can be as quick as 10 minutes [34], therefore it can be assumed that a significant proportion of the phage population are capable of ejecting their ICPs within 10 minutes of supplying the host.

It is hypothesised that the expelled, now cytoplasmic gp6.7::NtNL fusion could spontaneously complement with extracellular CtNL after immediately lysing the host cell, post-infection. Hypothetically, if following the same fate as wt-gp14, the gp14::NtNL fusion would form the outer pore of the ejectosome to aid with genomic DNA delivery into the host [20]. These membrane-lodged gp14::NtNL fusion proteins could hypothetically be exposed to the extracellular CtNL for immediate complementation, without the need for lysing the host cell.

Once extracellular NtNL::CtNL complementation is achieved, the next steps that are necessary to take in order to present this concept as a viable rapid bacterial detection system would be to calculate and fine tune the limit of detection (LoD) and speed of detection (SoD). In order to rival one of the most cutting edge, phage-based solutions [4], a LoD of 10^2^ CFU/mL and SoD of 3-7 hours would need to be achieved. Due to the fact that the ICP::fusion system bypasses host-TXTL and is constrained only by the time taken for the phage to adsorb to their host, it is expected that it has the potential to vastly outperform existing phage-based approaches, and potentially rival the fastest *E. coli* detection system that has been published [10]. However, for the ICP::fusion system to be able to match the sub-10 CFU/mL LoD displayed in state-of-the-art *E. coli* detection systems [7–10], it is expected that significant optimisations will need to be made.

Another point to consider is that, in its current state, the ICP::fusion system requires some form of hardware to measure the bioluminescent readout emitted when the host is detected. Moreover, whilst there have been significant developments in the field to increase the accessibility of measuring luminescence, including smartphone-based detection [35] and a commercial detection kit with the user-friendly EnSURE™ Touch hardware [36], it would be preferable to migrate to a colorimetric enzymatic system whereby a readout is produced that is visible to the naked eye. Fortunately, it is anticipated that only a simple alteration to the fusion construct design is necessary to facilitate this (i.e. replacing the NtNL sequence with an alternative enzyme sequence), and the overall motif and engineering approach of the system can remain unchanged.

To further explore the orthogonality of the diagnostic model displayed in this work, a similar engineering approach could be applied to the *Salmonella*-infecting phage P22 and Epsilon 15 - both of which have been shown to eject ICPs upon infection in a manner similar to T7 [37,38]. In doing so, this would demonstrate that the ICP::fusion system can detect three medically relevant pathogens: *E. coli K1* with engineered K1F phage, *Salmonella typhimurium* with engineered P22 phage, and *Salmonella anatum* with engineered Epsilon 15 phage. Nevertheless, regardless of the direction this proof-of-concept ICP::fusion system is propelled towards, it is anticipated that the key results displayed in this work have potential to play a significant role in the future of truly rapid and accessible bacterial diagnostics.

## 5. Patents

This section is not mandatory, but may be added if there are patents resulting from the work reported in this manuscript.

## Author Contributions

Conceptualization, J.P.W. and S.B.W.L.; methodology, J.P.W. and S.B.W.L.; validation, J.P.W., S.B.W.L. and T.F.; investigation, J.P.W.; resources, V.K. and A.P.S.; writing—original draft preparation, J.P.W.; writing—review and editing, J.P.W. and T.F.; visualization, J.P.W., S.B.W.L. and T.F.; supervision, V.K., A.P.S. and T.F.; project administration, V.K. and A.P.S.; funding acquisition, V.K. and A.P.S. All authors have read and agreed to the published version of the manuscript.

## Institutional Review Board Statement

Not applicable.

## Informed Consent Statement

Not applicable.

## Data Availability Statement

All the data are available upon request.

## Acknowledgments

In this section you can acknowledge any support given which is not covered by the author contribution or funding sections. This may include administrative and technical support, or donations in kind (e.g., materials used for experiments).

## Abbreviations

The following abbreviations are used in this manuscript:

AMR: Antimicrobial resistance
ELISA: Enzyme linked immunosorbent assay
AuNP: Gold nanoparticle
PCR: Polymerase chain reaction
NASBA: Nucleic acid sequence-based amplification
LAMP: Loop-mediated isothermal amplification
RPA: Recombinase polymerase amplification
GP: General practice
LFT: Lateral flow test
TXTL: transcription-translation
ICP: Internal capsid protein
CtNL: C-terminal sub-unit of NanoLuc
NtNL: N-terminal sub-unit of NanoLuc
Wt: Wild-type
PFU: Plaque forming units
RLU: Relative light units
GFP: Green fluorescent protein
LoD: Limit of detection
SoD: Speed of detection

## References

1. Zhang, D., Coronel-Aguilera, C., Romero, P., Perry, L., Minocha, U., Rosenfield, C., Gehring, A., Paoli, G., Bhunia, A. and Applegate, B. The Use of a Novel NanoLuc -Based Reporter Phage for the Detection of Escherichia coli O157:H7. Scientific Reports, 6(1), 2016.

2. Didovyk, A., Tonooka, T., Tsimring, L. and Hasty, J. Rapid and Scalable Preparation of Bacterial Lysates for Cell-Free Gene Expression. ACS Synthetic Biology, 6(12): 2198–2208, 2017.

3. Sun, Z., Hayes, C., Shin, J., Caschera, F., Murray, R. and Noireaux, V. Protocols for Implementing an Escherichia coli Based TX-TL Cell-Free Expression System for Synthetic Biology. Journal of Visualized Experiments, (79), 2013.

4. Wang, D., Chen, J. and Nugen, S. Electrochemical Detection of Escherichia coli from Aqueous Samples Using Engineered Phages. Analytical Chemistry, 89(3): 1650–1657, 2017.

5. Zhao, Y., Zeng, D., Yan, C., Chen, W., Ren, J., Jiang, Y., Jiang, L., Xue, F., Ji, D., Tang, F., Zhou, M. and Dai, J. Rapid and accurate detection of Escherichia coli O157:H7 in beef using microfluidic wax-printed paper-based ELISA. Analyst, 145(8): 3106–3115, 2020.

6. Zheng, L., Cai, G., Wang, S., Liao, M., Li, Y. and Lin, J. A microfluidic colorimetric biosensor for rapid detection of Escherichia coli O157:H7 using gold nanoparticle aggregation and smart phone imaging. Biosensors and Bioelectronics, 124-125: 143–149, 2019.

7. Kim, J. and Oh, S. Rapid detection of E. coli O157:H7 by a novel access with combination of improved sample preparation and real-time PCR. Food Science and Biotechnology, 29(8): 1149–1157, 2020.

8. Heijnen, L. and Medema, G. Method for rapid detection of viable Escherichia coli in water using real-time NASBA. Water Research, 43(12): 3124–3132, 2009.

9. Xia, X., Zhang, B., Wang, J., Li, B., He, K. and Zhang, X. Rapid Detection of Escherichia coli O157:H7 by Loop-Mediated Isothermal Amplification Coupled with a Lateral Flow Assay Targeting the z3276 Genetic Marker. Food Analytical Methods, vol. 15, 2021.

10. Hu, J., Wang, Y., Su, H., Ding, H., Sun, X., Gao, H., Geng, Y. and Wang, Z. Rapid analysis of Escherichia coli O157:H7 using isothermal recombinase polymerase amplification combined with triple-labeled nucleotide probes. Molecular and Cellular Probes, 50: 101501, 2020.

11. Kim, S. and Kim, S. Bacterial pathogen detection by conventional culture-based and recent alternative (polymerase chain reaction, isothermal amplification, enzyme linked immunosorbent assay, bacteriophage amplification, and gold nanoparticle aggregation) methods in food samples: A review. Journal of Food Safety, 41(1), 2020.

12. Deisingh, A. and Thompson, M. Strategies for the detection of Escherichia coli O157:H7 in foods. Journal of Applied Microbiology, 96(3):, 419–429, 2004.

13. Coleparmer.com. Rapid E. coli O157 and Shiga Toxin Antigen Detection Test Kits - Cole-Parmer. Available at https://www.coleparmer.com/p/rapid-e-coli-o157-and-shiga-toxin-antigen-detection-test-kits/15361 (accessed on 22 02 2022).

14. bioassayworks.com. E. coli O157:H7 Rapid Detection Kit. Available at: https://bioassayworks.com/?product=e-coli-o157h7-rapid-detection-kit-20-tests (accessed on 22 02 2022).

15. ,Shin, J. and Jardin, P. and Noireaux, V. Genome Replication, Synthesis, and Assembly of the Bacteriophage T7 in a Single Cell-Free Reaction. ACS Synthetic Biology, 1(9): 408–413, 2012.

16. Liyanagedera, S. and Williams, J. and Wheatley, J. and Biketova, A. and Hasan, M. and Sagona, A. and Purdy, K. and Puxty, R. and Feher, T. and Kulkarni, V. SpyPhage: A Cell-Free TXTL Platform for Rapid Engineering of Targeted Phage Therapies. ACS Synthetic Biology, 11 (10): 3330–3342, 2022.

17. Møller-Olsen, C., Ho, S., Shukla, R., Feher, T. and Sagona, A. Engineered K1F bacteriophages kill intracellular Escherichia coli K1 in human epithelial cells. Scientific Reports, 8(1), 2018.

18. Wisuthiphaet, N., Yang, X., Young, G. and Nitin, N. Rapid detection of Escherichia coli in beverages using genetically engineered bacteriophage T7. AMB Express, 9(1), 2019.

19. Kemp, P., Garcia, L. and Molineux, I. Changes in bacteriophage T7 virion structure at the initiation of infection. Virology, 340(2): 307–317, 2005.

20. Swanson, N., Hou, C. and Cingolani, G. Viral Ejection Proteins: Mosaically Conserved, Conformational Gymnasts. Microorganisms, 10(3), 2022.

21. Guo, F., Liu, Z., Fang, P., Zhang, Q., Wright, E., Wu, W., Zhang, C., Vago, F., Ren, Y., Jakana, J., Chiu, W., Serwer, P. and Jiang, W. Capsid expansion mechanism of bacteriophage T7 revealed by multistate atomic models derived from cryo-EM reconstructions. Proceedings of the National Academy of Sciences, 111(43): 4606–4614, 2014.

22. England, C., Ehlerding, E. and Cai, W. NanoLuc: A Small Luciferase Is Brightening Up the Field of Bioluminescence. Bioconjugate Chemistry, 27(5): 1175–1187, 2016.

23. Promega.co.uk. NanoLuc® Luciferase: One Enzyme, Endless Capabilities. Available at https://www.promega.co.uk/resources/technologies/NanoLuc-luciferase-enzyme/ (accessed on 28 02 2022).

24. Promega.com. NanoLuc®: A Smaller, Brighter, and More Versatile Luciferase Reporter. Available at https://www.promega.com/~/media/files/promega%20worldwide/europe/promega%20uk/webinars%20and%20events/cell%20analysis%20seminar%20tour/terry-riss-02.pdf (accessed on 28 02 2022).

25. Zhao, J., Nelson, T., Vu, Q., Truong, T. and Stains, C. Self-Assembling NanoLuc Luciferase Fragments as Probes for Protein Aggregation in Living Cells. ACS Chemical Biology, 11(1): 132–138, 2015.

26. Wheatley, J., Liyanagedera, S., Amaee, R., Sagona, A. and Kulkarni, V. Synthetic Biology for the Rapid, Precise and Compliant Detection of Microbes. In V. Singh (Ed.): Advances in Synthetic Biology, pp. 289–306, 2020.

27. Rustad, M. and Eastlund, A. and Jardine, P. and Noireaux, V. Cell-free TXTL synthesis of infectious bacteriophage T4 in a single test tube reaction. Synthetic Biology, 3 (1):, ysy002, 2018.

28. Lenk, E. and Casjens, S. and Weeks, J. and King, J. Intracellular visualization of precursor capsids in phage P22 mutant infected cells. Virology, /em 68 (1): 182–199, 1975.

29. Oliveira, L. and Tavaresa, P. and Alonso, C. Headful DNA packaging: Bacteriophage SPP1 as a model system. Virus Research, 173 (2): 247–259, 2013.

30. Hawkins, N. and Kizziah, J. and Penadés, J. and Dokland, T. Shape shifter: redirection of prolate phage capsid assembly by staphylococcal pathogenicity islands. Nature Communications, 12 (1): 5408, 2021,.

31. Cole, A. and Tran, S. and Ellington, A. Heat adaptation of phage T7 under an extended genetic code. Virus Evolution, 7 (2): veab100, 2021.

32. Swanson, N. and Lokareddy, R. and Li, F. and Hou, C. and Leptihn, S. and Pavlenok, M. and Niederweis, M. and Pumroy, R. and Moiseenkova-Bell, V. and Cingolani, G. Cryo-EM structure of the periplasmic tunnel of T7 DNA-ejectosome at 2.7 Å resolution. Molecular Cell, 81 (15): 3145–3159, 2021.

33. Molineux, I. No syringes please, ejection of phage T7 DNA from the virion is enzyme driven. Molecular microbiology, 40 (1), 2001.

34. You, L., Suthers, P. and Yin, J. Effects of Escherichia coli Physiology on Growth of Phage T7 In Vivo and In Silico. Journal of Bacteriology, 184(7), 1888–1894, 2002.

35. Kim, H., Jung, Y., Doh, I., Lozano-Mahecha, R., Applegate, B. and Bae, E. Smartphone-based low light detection for bioluminescence application. Scientific Reports, 7(1): 40203, 2017.

36. Hygiena. EnSURE™ Touch. Available at https://www.hygiena.com/food-safety-solutions/atp-monitoring/ensure-touch/ (accessed on 03 03 2022).

37. Wang, C., Tu, J., Liu, J. and Molineux, I. Structural dynamics of bacteriophage P22 infection initiation revealed by cryo-electron tomography. Nature Microbiology, 4(6): 1049–1056, 2019.

38. Chang, J., Schmid, M., Haase-Pettingell, C., Weigele, P., King, J. and Chiu, W. Visualizing the Structural Changes of Bacteriophage Epsilon15 and Its Salmonella Host during Infection. Journal of Molecular Biology, 402(4): 731–740, 2010.

